# ATX-2, the *C. elegans* Ortholog of Human Ataxin-2, Regulates Centrosome Size and Microtubule Dynamics

**DOI:** 10.1101/076604

**Authors:** Michael D. Stubenvoll, Jeffrey C. Medley, Miranda Irwin, Mi Hye Song

## Abstract

Centrosomes are critical sites for orchestrating microtubule dynamics, and exhibit dynamic changes in size during the cell cycle. As cells progress to mitosis, centrosomes recruit more microtubules (MT) to form mitotic bipolar spindles that ensure proper chromosome segregation. We report a new role for ATX-2, a *C. elegans* ortholog of Human Ataxin-2, in regulating centrosome size and MT dynamics. ATX-2, an RNA-binding protein, forms a complex with SZY-20 in an RNA-independent fashion. Depleting ATX-2 results in embryonic lethality and cytokinesis failure, and restores centrosome duplication to *zyg-1* mutants. In this pathway, SZY-20 promotes ATX-2 abundance, which inversely correlates with centrosome size. Centrosomes depleted of ATX-2 exhibit elevated levels of centrosome factors (ZYG-1, SPD-5, γ-Tubulin), increasing MT nucleating activity but impeding MT growth. We show that ATX-2 influences MT behavior through γ-Tubulin at the centrosome. Our data suggest that RNA-binding proteins play an active role in controlling MT dynamics and provide insight into the control of proper centrosome size and MT dynamics.

## Introduction

As the primary microtubule-organizing centers, centrosomes are vital for the maintenance of genomic integrity in animal cells (Nigg and Stearns 2011). The centrosome consists of two barrel-shaped centrioles surrounded by a network of proteins termed pericentriolar materials (PCM). To maintain the fidelity of cell division, each cell must duplicate a pair of centrioles precisely once per cell cycle, one daughter per mother centriole. Mishaps in centrosome assembly result in chromosome missegregation and other cell cycle defects. Thus, stringent regulation of centrosome assembly is imperative for proper cell division and survival.

Studies in *C. elegans* have discovered five evolutionarily conserved proteins (ZYG-1, SPD-2, SAS-4, SAS-5 and SAS-6) that are required for centrosome assembly (Delattre et al., 2006; Habedanck et al., 2005; Pelletier et al., 2006; Shimanovskaya et al., 2014). Many other factors, including protein phosphatase 2A, also regulate the production, activity, or turnover of core regulators, and are equally important in regulating centrosome assembly (Kitagawa et al., 2011; Song et al., 2011). Like other biological processes, centrosome assembly is regulated by a combined action among negative and positive regulators (Kemp et al., 2007). While the kinase ZYG-1 promotes centriole duplication in *C. elegans*, *szy-20* acts as a genetic suppressor of *zyg-1* (O’Connell et al., 2001; Song et al., 2008). The *szy-20* gene encodes a centrosome-associated RNA-binding protein that negatively regulates centrosome assembly by opposing ZYG-1. Centrosomes in *szy-20* mutants exhibit elevated levels of centrosomal proteins, resulting in defective microtubule (MT) behavior and embryonic lethality. SZY-20 contains putative RNA-binding domains (SUZ, SUZ-C). Mutating these domains has been shown to perturb *in vitro* RNA-binding of SZY-20 and its capacity to regulate centrosome size *in vivo* (Song et al., 2008). Other studies have shown that a number of RNAs and RNA-binding proteins are associated with centrosomes and MTs, and influence proper mitotic spindles and other aspects of cell division. In mammalian cells, several RNA-binding proteins (e.g., RBM8A, Hu antigen R, and the Ewing sarcoma protein) associate with centrosomes and play a role in regulating centrosome assembly during cell division (Castro et al., 2000; Filippova et al., 2012; Ishigaki et al., 2014; Leemann-Zakaryan et al., 2009; Ugrinova et al., 2007). In yeast, spindle pole body duplication is linked to translational control via the action of RNA-binding proteins (Sezen et al., 2009). In *Xenopus*, MT-guided localization of transcripts followed by spatially enriched translation is important for proper MT behavior and cell division, suggesting the importance of local translational control (Blower et al., 2005; Blower et al., 2007; Groisman et al., 2000).

Despite the finding that SZY-20 negatively regulates ZYG-1, no direct interaction between the two proteins has been found. Thus, identifying additional factors that function between SZY-20 and ZYG-1 should provide further insights into the molecular mechanism by which the putative RNA-binding protein, SZY-20, influences centrosome assembly. Toward this end, we report here our identification of an RNA-binding protein ATX-2 that physically associates with SZY-20. ATX-2 is the *C. elegans* ortholog of human Ataxin-2 that is implicated in human neurodegenerative disease (Lastres-Becker et al., 2008). Specifically, human spinocerebellar ataxia type 2 is shown to be associated with an extended poly-glutamine (Q) tract in Ataxin-2 (Elden et al., 2010; Huynh et al., 1999; Lastres-Becker et al., 2008). Ataxin-2 is an evolutionarily conserved protein that contains an RNA-binding motif (LSm: Sm-like domain) and a PAM domain for binding the poly-(A) binding protein (PABP1) (Achsel et al., 2001; Jimenez-Lopez and Guzman 2014; Satterfield and Pallanck, 2006). It has been shown that Ataxin-2 binds directly to the 3’UTR of mRNAs and stabilize target transcripts, and that poly-Q expansion blocks the RNA-binding by Ataxin-2 *in vitro* (Yokoshi et al., 2014). Ataxin-2 homologs have been implicated in a wide range of RNA metabolism-dependent processes including translational control of circadian rhythm (Zhang et al., 2013; Lim and Allada, 2013). While ATX-2 in *C. elegans* is known to be responsible for embryonic development and translational control in germline development (Ciosk et al., 2004; Kiehl et al., 2000; Maine et al., 2004; Skop et al., 2004), the action of this RNA-binding protein in centrosome assembly and cell division has not been fully explored.

In this study, we investigate the role of *C. elegans* ATX-2 in early cell cycles and how ATX-2 acts together with SZY-20 and ZYG-1 in controlling centrosome size and MT behavior. We show that ATX-2 negatively regulates the key centriole factor ZYG-1. In the centrosome assembly pathway, SZY-20 acts upstream of ATX-2 and positively regulates embryonic levels of ATX-2; proper levels of ATX-2 contributes in turn to normal centrosome size and subsequent MT dynamics.

## Results

### ATX-2 physically interacts with SZY-20 *in vivo* independently of RNA

To further elucidate the role of SZY-20 in regulating centrosome assembly, we looked for additional factors interacting with SZY-20. By immunoprecipitating (IP) endogenous SZY-20 from worm protein lysates with anti-SZY-20, followed by mass spectrometry, we generated a list of proteins associated with SZY-20 *in vivo*. As for a putative RNA-binding protein SZY-20, we found many known RNA-binding proteins co-precipitated with SZY-20, including ATX-2 (11 peptides, 19% coverage = 181/959aa) and PAB-1 (15 peptides, 32% coverage = 206/646aa) (**S1A Fig**). In *C. elegans*, ATX-2 is shown to form a cytoplasmic complex with PAB-1 (Ciosk et al., 2004), and the ATX-2-PAB-1 interaction is conserved from yeast to mammals (Ciosk et al., 2004; Nonhoff et al., 2007; Satterfield and Pallanck, 2006; Yokoshi et al., 2014). *pab-1* encodes Poly-A-binding protein, a *C. elegans* homolog of a human PABP1 that plays a role in RNA stability and protein translation (Ciosk et al, 2004; Kahvejian et al., 2005).

To confirm the physical interaction between SZY-20 and ATX-2, we used anti-SZY-20 to pull down SZY-20 and its associated proteins from embryonic lysates and examined co-precipitates by western blot (**Fig 1A**). Consistent with our mass spectrometry data, we detected ATX-2 and SZY-20 in the SZY-20-immunoprecipitates from wild-type embryonic extracts. Given that this protein complex consists of RNA-binding proteins, we asked if the physical association is mediated through RNA. To test RNA dependence, we repeated IP to pull down SZY-20 interacting proteins in the presence of RNaseA or RNase inhibitor and found that they co-precipitated in either condition (**Fig 1A, S1B Fig**), suggesting ATX-2 and SZY-20 physically interact in an RNA-independent manner although we cannot exclude the possibility that RNA bound by the ATX-2-SZY-20 complex could have been protected from RNase treatment.

**Fig. 1.**
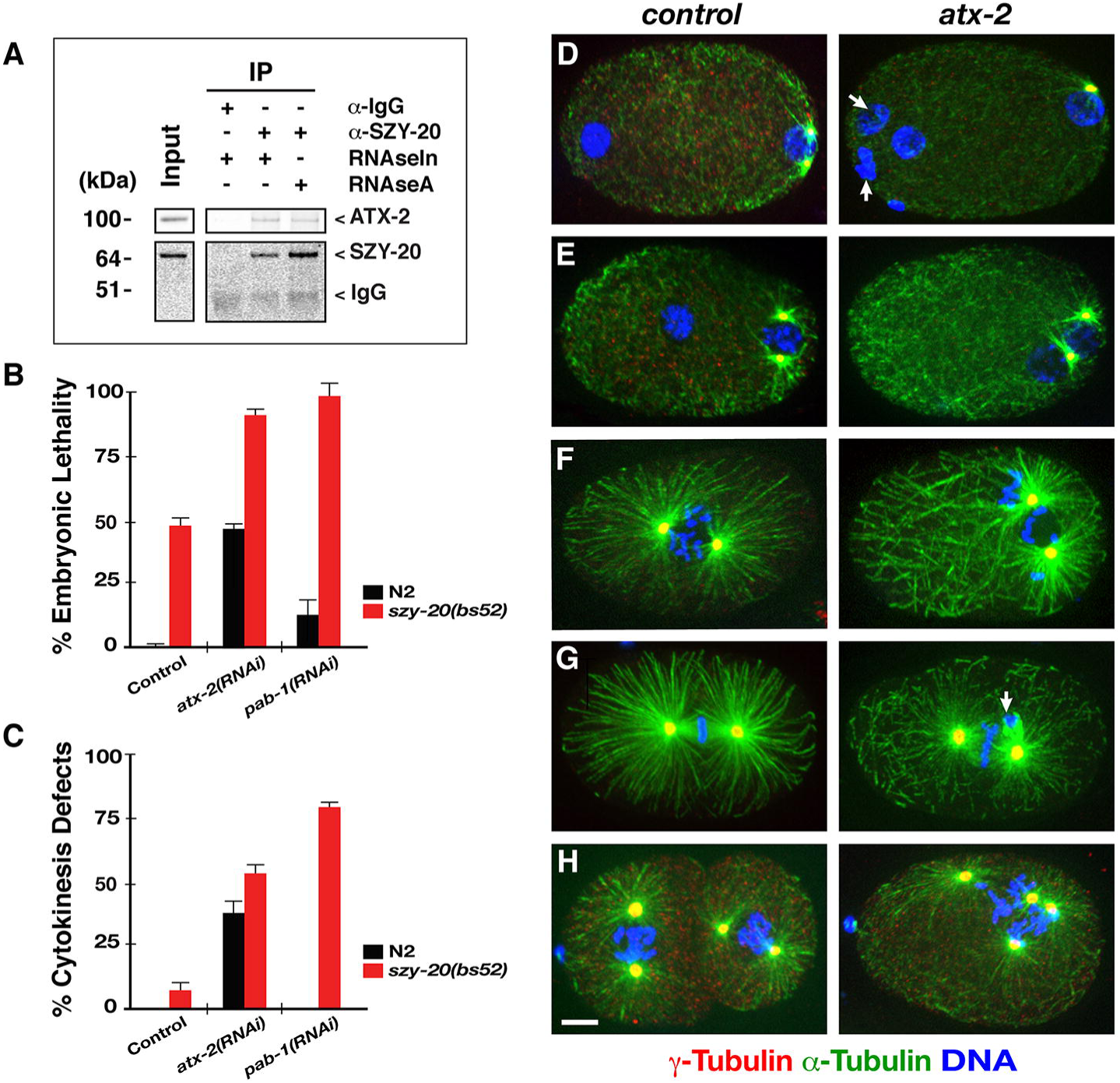
ATX-2 physically associates with SZY-20 and functions in early cell division. (A) ATX-2 co-precipitates with SZY-20 in the presence of RNAse inhibitor or RNAse A. For input, 10% of the total embryonic lysates was loaded for comparison. (B) At 24°C, *atx-2(RNAi)* and *szy-20(bs52)* animals produce 50-60% of embryonic lethality but *pab-1(RNAi)* produces < 10% of embryonic lethality. Depleting *atx-2* or *pab-1* in *szy-20(bs52)* mutants leads to a significant increase in embryonic lethality (**Table 1**, n=500-1500). (C) Synergistic effect on abnormal cytokinesis by co-depleting *szy-20* with *atx-2* or *pab-1*: At 24°C, 6% of *szy-20(bs52)* but none (n=68) of *pab-1(RNAi)* or 36% (n=114) of *atx-2(RNAi)* embryos fail to complete cytokinesis. However, *atx-2(RNAi)* and *pab-1(RNAi)* in *szy-20(bs52)* mutant embryos produce 54% (n=86) and 82% (n=68) of cytokinesis failure, respectively. (D-H) Compared to control (L4440), *atx-2(RNAi)* results in cell division defects such as (D) polar body extrusion failure marked with extra DNA (arrows, 22%, n=114; **S3 Movie**), (E-F) abnormal spindle positioning (3%, n=68), (G) chromosome missegregation (arrow, 10%, n=84), (H) cytokinesis failure (36%, n=114) marked by extra DNA and supernumerary centrosomes in single-cell embryos. Note short spindle and astral MTs in *atx-2(RNAi)* embryos. *atx-2(ne4297)* embryos exhibit similar cell cycle defects (**Fig 3B**). Error bars are the standard deviation (SD). Bar, 5 μm.

The *szy-20(bs52)* mutation results in a truncated protein, deleting the C-terminal 197 aa residues including the SUZ-C domain, one of the putative RNA-binding domains in SZY-20 (Song et al., 2008). We utilized *szy-20(bs52)* embryos for IP analysis to determine if the C-terminal truncation of SZY-20 affected physical association of SZY-20 with ATX-2 (**S1B Fig**).

Whereas ATX-2 co-precipitated with SZY-20 in wild-type extracts, ATX-2 was undetectable in co-precipitates from *szy-20(bs52)* extracts, suggesting that the C-terminus of SZY-20 influences physical interaction with ATX-2. IP assay using embryos expressing SZY-20-GFP-3xFLAG yielded a similar result: ATX-2 is undetectable in co-precipitates of SZY-20 tagged with GFP-3xFLAG at the C-terminus (**S1E Fig**), supporting that proper folding of the C-terminal domain is critical for SZY-20 to interact with ATX-2. Further, the C-terminal deletion in either ATX-2 or SZY-20 appears to alleviate its interaction with SZY-20 (**S1C and S1D Fig**). By additional IP assays with anti-GFP using embryos expressing various GFP-tagged proteins, we further confirmed that both ATX-2 and PAB-1 physically interact with SZY-20 or with each other (**S1E Fig)**. Together, our data suggest SZY-20 forms a complex with known RNA-binding proteins ATX-2 and PAB-1 *in vivo,* via direct or indirect interaction.

### ATX-2 is required for proper cell division during embryogenesis

We next asked if these SZY-20 interacting proteins play a similar role to SZY-20 during early cell division. In *C. elegans,* ATX-2 is required for translational control in gonadogenesis and for normal cytokinesis during embryogenesis (Ciosk et al., 2004; Kiehl et al., 2000; Maine et al., 2004; Skop et al., 2004). To date, PAB-1 is only known to be required for germline development (Ciosk et al., 2004; Ko et al., 2013). Consistent with prior studies, we observed strong embryonic lethality in both *atx-2(RNAi)* and *atx-2(ne4297)* but a very low embryonic lethality in *pab-1(RNAi)*, and sterility by loss of *pab-1* or *atx-2* (**Fig 1B**).

Embryonic lethality by loss of *atx-2* might result from defective cell division. To examine what role ATX-2 plays in cell division, we immunostained embryos for microtubules (MTs), centrosomes and DNA (**Fig 1D-1H**). Confocal microscopy of immunofluorescence (IF) revealed that knocking down *atx-2* by RNAi results in multiple cell division defects including polar body extrusion failure (22%; **S3 Movie**), abnormal spindle positioning (3%), chromosome missegregation (10%) and cytokinesis failure (36%; n=114). We observed similar cell division phenotypes, but with higher penetrance in temperature sensitive (ts) *atx-2(ne4297)* mutants. By 4D time-lapse confocal microscopy, we observed that incomplete cytokinesis following successful centrosome duplication results in tetrapolar spindles in one-cell embryo (**S1 and S2 Movie**). In these embryos, the cytokinetic furrow initiates but cytokinesis fails to complete, resulting in a multi-nucleated cell with four centrosomes after the second mitosis. All of these cell division phenotypes resemble cell cycle defects observed previously in *szy-20(bs52)* embryos (Song et al, 2008), suggesting that ATX-2 functions closely with SZY-20 in cell division. In contrast, *pab-1(RNAi)* produced only minor cell cycle defects such as atypical spindle positioning (**S2A Fig**).

Since ATX-2 appears to play a similar role to SZY-20, we then asked if *atx-2* genetically interacts with *szy-20*. Genetic analysis by combining *atx-2(RNAi)* or *pab-1(RNAi)* with a hypomorphic mutant allele *szy-20(bs52)* suggest a positive genetic interaction among these factors (**Fig 1B, Table 1**). At 24°C, the semi-restrictive temperature, *szy-20(bs52)* animals fed with *atx-2(RNAi)* produced higher embryonic lethality than single knockdown of either *szy-20(bs52)* or *atx-2(RNAi)*. Similarly, *pab-1(RNAi)* in *szy-20(bs52)* led to a synergistic increase in embryonic lethality compared to *szy-20(bs52)* or *pab-1(RNAi)* alone.

**Table 1.** Genetic Analysis of atx-2.

| | °C | % Embryonic Lethality<br>(ave ± s.d.) | N<sup>a</sup> |
| --- | --- | --- | --- |
| *N2* | 24 | 1 ± 1.0 | 501 |
| *szy-20(bs52)* | | 49 ± 4.2 | 523 |
| *atx-2(RNAi)* | | 47 ± 3.4 | 1499 |
| *atx-2(ne4297)* | | 100 ± 0.3 | 909 |
| *pab-1(RNAi)* | | 9 ± 7.0 | 584 |
| *szy-20(bs52); atx-2(RNAi)* | | 91 ± 4.0 | 821 |
| *szy-20(bs52); atx-2(ne4297)* |  | sterile | - |
| *pab-1(RNAi); szy-20(bs52)* | | 97 ± 6.0 | 657 |
| *N2* | 22 | 0 ± 1.0 | 668 |
| *zyg-1(it25)* | | 98 ± 1.0 | 544 |
| *atx-2(ne4297)* | | 99 ± 1.0 | 361 |
| *zyg-1(it25); atx-2(ne4297)* | | 98 ± 3.0 | 304 |
| *N2* | 20 | 0 ± 0.0 | 1053 |
| *zyg-1(it25)* | | 1 ± 2.0 | 1107 |
| *atx-2(ne4297)* | | 74 ± 18.0<sup>b</sup> | 937 |
| *zyg-1(it25); atx-2(ne4297)* | | 63 ± 24.0<sup>b</sup> | 1057 |
| *N2* | 18 | 0 ± 0.0 | 747 |
| *zyg-1(it25)* | | 0 ± 0.0 | 705 |
| *atx-2(ne4297)* | | 45 ± 7.0<sup>c</sup> | 324 |
| *zyg-1(it25); atx-2(ne4297)* | | 32 ± 12.0<sup>c</sup> | 383 |

<sup>a</sup> The number of progeny scored.
<sup>b</sup> *P*-value=0.2 (Student *t*-test, two-tailed distribution and two-sample equal variance)
<sup>c</sup> *P*-value=0.02 (Student *t*-test, two-tailed distribution and two-sample equal variance)

Cytological analyses further confirmed the functional interaction among these factors. Double-knockdown significantly enhanced the penetrance of cell division phenotypes, compared to mock-treated *szy-20(bs52)* or RNAi alone: 55% (n=128) of *szy-20(bs52); atx-2(RNAi)* embryos exhibit a failure of polar body extrusion, compared to 22% (n=114) of *atx-2(RNAi)* and 18% (n=145) of *szy-20(bs52)*. This synergistic effect is even stronger for cytokinesis: 36% of *atx-2(RNAi)* and 6% of *szy-20(bs52),* but 68% of *szy-20(bs52); atx-2(RNAi)* embryos failed to complete cytokinesis at first division (**Fig 1C, 1H)**. Consistent with embryonic lethality, *pab-1(RNAi); szy-20(bs52)* produced highly penetrant cytokinesis defect, in contrast to the weak phenotype by *pab-1(RNAi)* alone (**Fig 1C)**. Combining *pab-1(RNAi)* and *szy-20(bs52)* mutation produced 82% (n=68) of cytokinesis failure and 24% (n=68) of polar body extrusion failure, while *pab-1(RNAi)* (n=68) alone showed a minor cell cycle defect (**S2A Fig**). In addition, *atx-2(RNAi)* or *pab-1(RNAi)* enhances sterility in *szy-20(bs52)* animals, consistent with the sterility associated with all three genes. Together, our data suggest that RNA-binding proteins ATX-2 and PAB-1 function in close association with SZY-20 during embryogenesis.

### ATX-2 negatively regulates centrosome assembly

As *szy-20* is known as a genetic suppressor of *zyg-1* (Song et al., 2008), we asked if ATX-2/PAB-1 functions in centrosome assembly as well. To address this, we first examined centrosome duplication in *zyg-1(it25)* embryos fed with *atx-2* or *pab-1(RNAi)* at 24°C (**Fig 2**). When grown at 24°C the restrictive temperature, *zyg-1(it25*) embryos fail to duplicate centrosomes during the first cell cycle, producing monopolar spindles at the second cell division (O’Connell et al., 2001). By recording live imaging of *zyg-1(it25)* embryos expressing GFP-α-Tubulin, mCherry-γ-Tubulin and mCherry-Histone, we scored centrosome duplication at the second mitosis (**Fig 2A-2C**). Over 60% of *atx-2* or *pab-1(RNAi)* treated *zyg-1(it25)* embryos produced bipolar spindles, indicating successful centrosome duplication during the first cell cycle, while only 3% of mock treated *zyg-1(it25)* embryos formed bipolar spindles. We noticed, however, that a great majority of *atx-2(RNAi)* treated *zyg-1(it25)* embryos exhibit four centrosomes in one-cell embryo after the second cell cycle, suggestive of cytokinesis failure due to loss of *atx-2* (**Fig 2C, S1 and S2 Movie**). Consistent with RNAi-mediated knockdown, we made a similar observation in *zyg-1(it25); atx-2(ne4297)* double mutants (**S3A and S3B Fig**).

**Fig 2.**
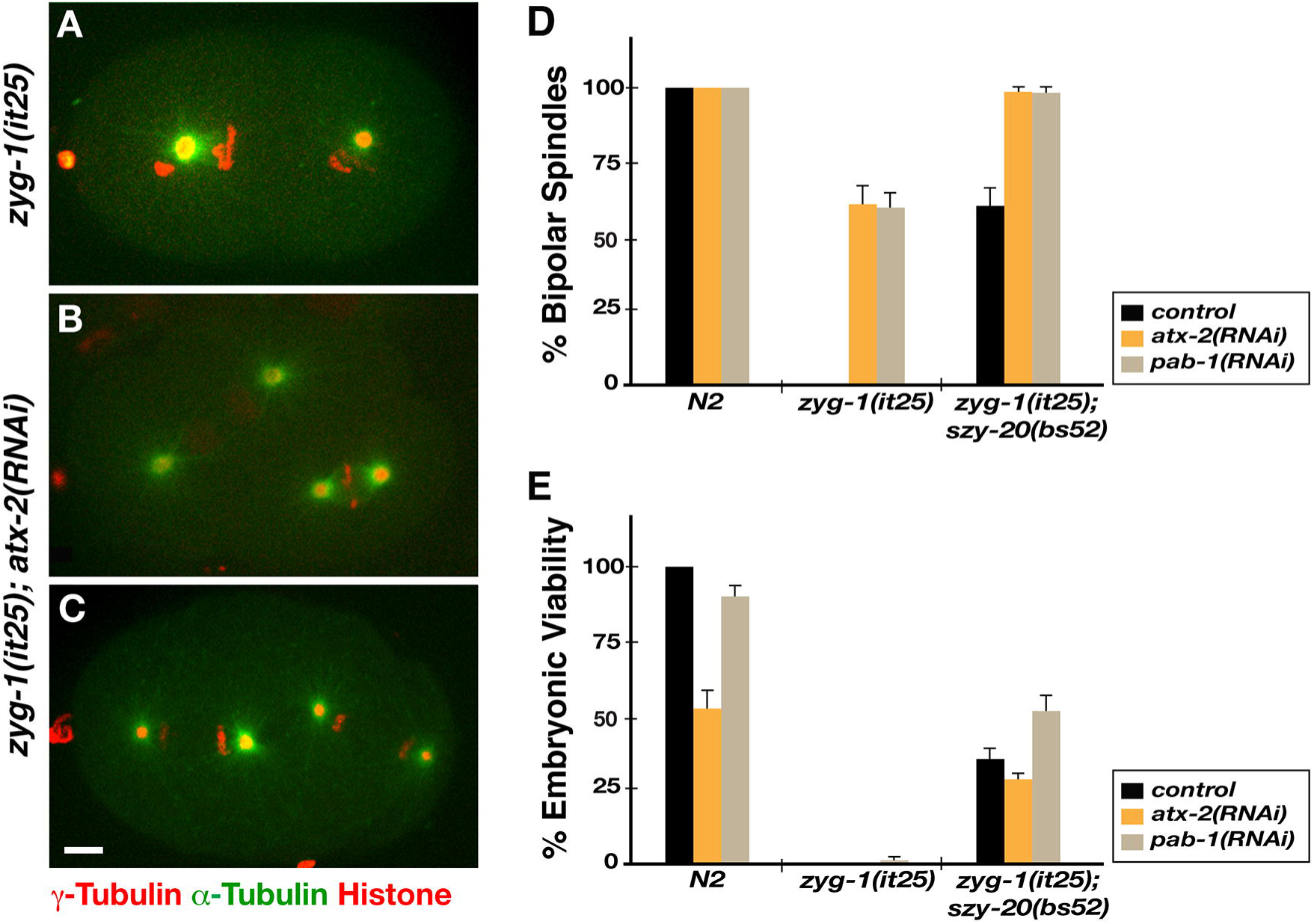
ATX-2 negatively regulates centrosome assembly. (A) *zyg-1(it25)* treated with control *RNAi* (*L4440*) produces monopolar spindles at the second cell cycle. (B-C) *atx-2(RNAi)* restores bipolar spindle formation to *zyg-1(it25)* embryos grown at 24°C. (C) Note that *zyg-1(it25); atx-2(RNAi)* embryo exhibits four spindle poles in one-cell embryo at the second cell cycle, indicative of cytokinesis failure caused by loss of *atx-2*. Bar, 5 μm. (D) Quantification of bipolar spindle formation at 24°C. While *atx-2(RNAi)* or *pab-1(RNAi)* in *N2* (wild-type) has no effect on centrosome duplication, either RNAi restores >60% of bipolar spindles to *zyg-1(it25)* embryos (n > 200) at the second mitosis, suggesting restoration of centrosome duplication during the first cell cycle. Co-depletion with *szy*-20 further enhances % bipolar spindles to *zyg-1(it25)*: 97% of *atx-2(RNAi)* (n=66) or 98% of *pab-1(RNAi)* (n=64) in *zyg-1(it25); szy-20(bs52)* produces bipolar spindles, compared to ~60% of bipolar spindles in single knockdown in *szy-20* (n=116) or others. (E) At the semi-restrictive temperature 23°C, *atx-2* or *pab-1(RNAi)* partially restores embryonic viability to *zyg-1(it25),* when co-depleted with *szy-20*. *atx-2* (27%, n=120) or *pab-1(RNAi)* (52%, n=139) with *szy-20(bs52)* mutation leads to embryonic viability to *zyg-1(it25)*. Single-knockdown of *atx-2* or *pab-1* show no significant effect on embryonic viability in *zyg-1(it25).* Error bars are SD.

Given the positive genetic interactions among *szy-20, atx-2* and *pab-1*, we further asked if co-depleting these factors could enhance the suppression of *zyg-1* (**Fig 2D, 2E**). At 24°C, co-depleting either *atx-2* or *pab-1* with *szy-20* restored nearly 100% of centrosome duplication to *zyg-1(it25)* embryos, while single depletion produced ~60% duplication in *zyg-1(it25)*. Despite the restoration in centrosome duplication at 24°C, none of these embryos hatched, owing to other cell cycle defects such as cytokinesis failure described above. However, we were able to show partial restoration of embryonic viability in *zyg-1(it25)* by co-depleting either *atx-2* or *pab-1* with *szy-20* at semi-restrictive temperature 23°C, whereas single depletion of *atx-2* or *pab-1* showed no effect on embryonic viability of *zyg-1(it25).* Our data indicate that like *szy-20*, *atx-2* and *pab-1* act as genetic suppressors of *zyg-1*. Thus, these RNA-binding proteins in a complex function together to negatively regulate centrosome assembly.

Finally, we asked if *any* RNA-binding protein could suppress *zyg-1* through global translational control. We chose the RNA-binding protein CAR-1 that is required for cytokinesis (Audhya et al., 2005; Squirrell et al., 2006) to test if *car-1* suppresses *zyg-1* (**S3C Fig**). We found no sign of centrosome duplication in *car-1(RNAi); zyg-1(it25)* embryos, while *atx-2(RNAi)* restored centrosome duplication to *zyg-1(it25)* in a parallel experiment. This result suggests that RNA-binding proteins are not general suppressors of *zyg-1*, but that actions of RNA-binding proteins (SZY-20/ATX-2/PAB-1) are specific to the ZYG-1 dependent centrosome assembly pathway.

### SZY-20 promotes ATX-2 abundance in early embryos

To understand how these RNA-binding proteins coordinate their actions within the cell, we examined the subcellular localization in early embryos. Immunostaining embryos with α-ATX-2 shows a diffuse pattern throughout the cytoplasm with occasional small foci; as expected, this signal is significantly diminished in *atx-2(RNAi)* or *atx-2(ne4297)* embryos (**Fig 3A, 3B**). We also generated a transgenic strain expressing *atx-2-gfp-3x flag* (**S1 Table**) and observed the dynamics of ATX-2-GFP using time-lapse movies, which shows consistent patterns to endogenous ATX-2 (**S4 Movie**). *C. elegans* PAB-1 is also found in the cytoplasm and is often associated with P-bodies or stress granules (Gallo et al., 2008) and SZY-20 localizes to the nuclei, centrosome and cytoplasm in early embryos (Song et al., 2008). Co-staining embryos show that ATX-2 partially coincides with cytoplasmic SZY-20 (**Fig 3C, S2B Fig)** or PAB-1 (**Fig 3D, S2C Fig**). A similar co-localization is also observed for GFP-PAB-1 and SZY-20 (**S2D Fig**). Thus, these RNA-binding proteins appear to function together in the cytoplasm to regulate early cell division.

**Fig 3.**
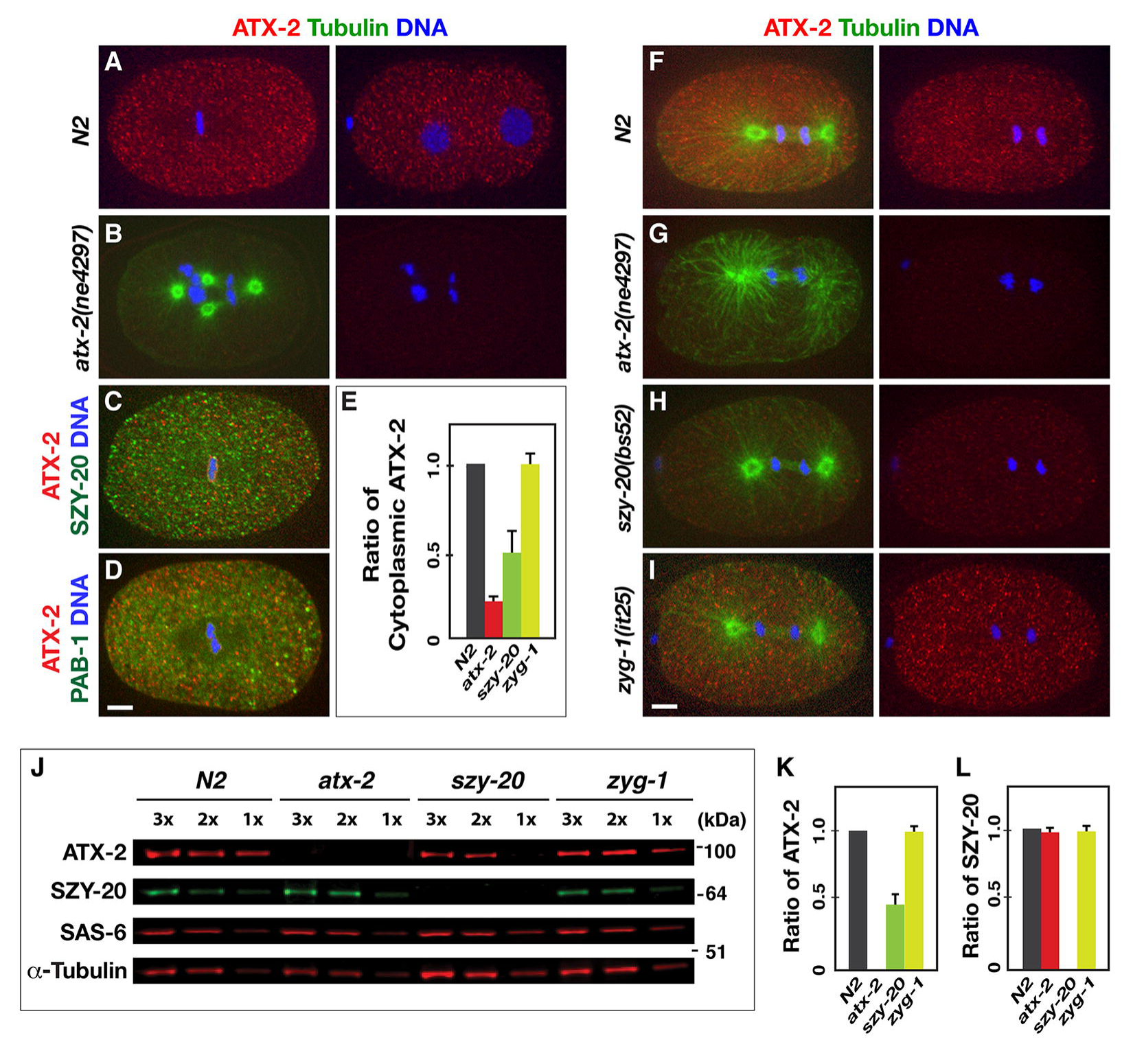
Subcellular localization and molecular hierarchy of ATX-2. Immunostaining illustrates (A) a diffuse distribution of ATX-2 in the cytoplasm in one-cell (left) and two-cell (right) stage embryos, and (B) a reduced staining in *atx-2(ne4297)* embryo that exhibits cytokinesis failure. (C) Wild-type embryo co-stained for ATX-2 and SZY-20 illustrates that ATX-2 and SZY-20 partially coincide in the cytoplasm (**S2B Fig**). (D) Wild-type embryo stained for ATX-2 and GFP-PAB-1 illustrates that ATX-2 and PAB-1 partially coincide in the cytoplasm (**S2C Fig**). (E-I) Quantification (E) of cytoplasmic ATX-2 levels reveals that *szy-20(bs52)* embryos possess ~50% (n=22) of normal ATX-2 levels. *atx-2(ne4297)* embryos possess ~18% (± 9, n=18) of cytoplasmic ATX-2, while *zyg-1(it25)* embryos exhibit a normal level (96% ± 6, n=17) of ATX-2, compared to N2 embryos (n=15). (F-I) Embryos at first mitosis show that cytoplasmic expression of ATX-2 is affected by genetic mutations compared to N2°Control. *atx-2(ne4297)* embryo exhibits a great reduction in ATX-2 staining (G). ATX-2 cytoplasmic staining is significantly reduced in *szy-20(bs52)* embryo (H), but no changes in *zyg-1(it25)* embryo (I). Shown are images from a single plane of the embryo. Bar, 5 μm. (J-L) Immunoblot using embryonic lysates from *atx-2, szy-20, zyg-1* mutants and N2 worms grown at 22°C. γ-Tubulin was used as loading control. (K-L) Quantitative immunoblot analyses reveal (K) *szy-20(bs52)* embryos possess significantly reduced levels of ATX-2 (0.4 ± 0.1, n=10) compared to N2 (n=12), (L) whereas *atx-2(ne4297)* embryos contain normal levels of SZY-20 (0.98 ± 0.03, n=12). Note that full-length protein levels are quantified for ATX-2 and SZY-20, as both *atx-2(ne4297)* and *szy-20(bs52)* embryos produce a truncated polypeptide. In contrast, no significant changes were found in the level of a centriole factor, SAS-6, (0.97 ± 0.1, n=12) in any mutant embryos. Error bars are SD.

To gain insights into the hierarchical relationship among these factors, we used mutant strains to determine how the embryonic expression is affected by the other factors (**Fig 3E-3I**). Quantitative immunofluorescence revealed that ATX-2 levels in the cytoplasm are significantly reduced (~50%) in the hypomorphic *szy-20(bs52)* embryo relative to wild-type control (**Fig 3E, 3H**), while the *zyg-1(it25)* mutation had no significant effect on the cytoplasmic ATX-2 (**Fig 3E, 3I**). SZY-20 localization was, however, unaffected by either *atx-2* or *pab-1* depletion (**S2D Fig**). Consistently, prior study showed that *szy-20* acts upstream of *zyg-1* (Song et al., 2008). Thus, it seems likely that *szy-20* function upstream of *atx*-2, with both acting upstream of *zyg-1.* Quantitative western blots further support this hierarchical relationship (**Fig 3J-3L**). *szy-20(bs52)* embryos exhibit significantly reduced ATX-2 levels (~40%; **Fig 3K**) relative to wild-type control, while *atx-2(ne4297)* embryos show normal levels of SZY-20 (~98%; **Fig 3L**) and *zyg-1* mutants contain normal levels of both ATX-2 and SZY-20 (**Fig 3J-3L**). However, no significant changes are found in the level of a centriole factor, SAS-6 in either mutant embryo (**Fig 3J**). Taken together, our data suggest that SZY-20 acts upstream of ATX-2, and positively regulates ATX-2 abundance in early embryos.

### Loss of *atx-2* leads to increased levels of centrosomal ZYG-1

Because *atx-2* acts as a genetic suppressor of *zyg-1,* we reasoned that inhibiting ATX-2 might enhance ZYG-1 activity, thereby restoring centrosome duplication and embryonic viability to *zyg-1(it25)* embryos. By staining embryos for ZYG-1 and microtubules (**Fig 4A**), we quantified the fluorescence intensity of ZYG-1 at first metaphase centrosomes, finding that *atx-2* mutant centrosomes possess twice as much ZYG-1 levels as those in control embryos (*p <*0.001, **Fig 4B**). Using the CRISPR-Cas9 method (Dickinson et al., 2015; Paix et al., 2015), we also generated a strain expressing HA-tagged ZYG-1 at endogenous levels from the native genomic locus **(S4A-S4C Fig, S2 Table)**. By labeling endogenous ZYG-1 with anti-HA **(S4A and S4B Fig**), we observed a similar pattern of ZYG-1 localization at centrosomes to that seen by anti-ZYG-1 **(S4E Fig**). The levels of centrosomal HA-ZYG-1 are also increased by two-fold in *atx-2(RNAi)* embryos **(S4C Fig)**.

**Fig 4.**
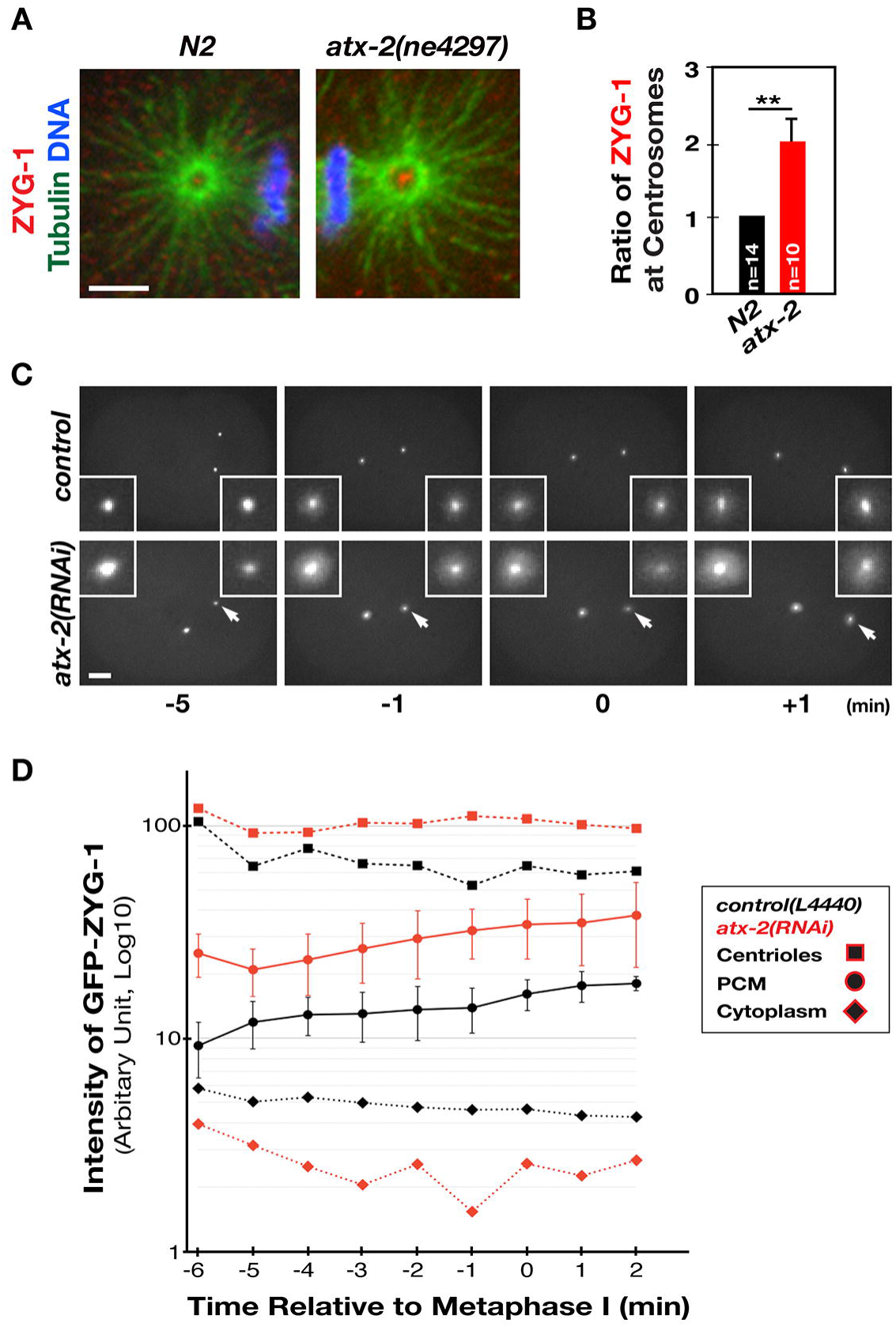
Loss of *atx-2* results in increased levels of ZYG-1 at centrosomes. (A) The *atx-2*mutant centrosome exhibits more intense ZYG-1 focus than N2. (B) Relative fluorescence intensity of centrosomal ZYG-1 at the first metaphase (\*\**p<*0.001). (C) Still images from time-lapse movies of embryos expressing GFP-ZYG-1-C-term (Peters et al., 2010) at selected time points. Time (min) is relative to the first metaphase. These embryos exhibit centriolar as well as pericentriolar GFP-ZYG-1, with increased levels in the *atx-2(RNAi)* embryo. In the *atx-2(RNAi)* embryo, the centrosome on the right (arrow) is out of the focal plane as two centrosomes in *atx-2(RNAi)* embryos often do not align on the same focal plane (see **S5 and S6 Movie**). (D) Measurements of fluorescence intensity of centriolar, pericentriolar and cytoplasmic GFP-ZYG-1 using time-lapse recordings (n=5-7). ZYG-1 levels steadily increase in control and *atx-2(RNAi)* embryos as cell cycle progresses. Note a slightly lower cytoplasmic levels of ZYG-1 in *atx-2(RNAi)* embryos. Error bars are SD (centriolar and pericentriolar ZYG-1; *p*<0.0001; cytoplasmic ZYG-1; *p*>0.005). Bar, 5 μm.

Given that ZYG-1 localizes to centrosomes in a cell cycle-dependent manner, the observed increase in centrosomal ZYG-1 at first metaphase in *atx-2* mutants could result from a shift in the cell cycle due to loss of *atx-2*. To examine cell cycle dependence of ZYG-1 localization to centrosomes, we utilized a strain expressing GFP-ZYG-1-C-term that contains a C-terminal portion (217-706 aa) lacking most of the kinase domain, but including the Cryptic Polo Box (CPB) that is sufficient for centrosomal targeting (Peters et al., 2010; Shimanovskaya et al., 2014). To observe dynamics of centrosome-associated ZYG-1 over time, we acquired 4D time-lapse movies of early embryos starting from pronuclear meeting up to separation of the centriole pair at first anaphase (**Fig 4C, 4D, S5 and S6 Movie**). Using these recordings, we first quantified the fluorescence intensity of the intensely labeled sub-centrosomal GFP signal, which presumably reflects the centriolar structure. Then, we measured pericentriolar GFP signal that likely represents PCM. Throughout the cell cycle, we observe a nearly two-fold increase in both centriolar and PCM-associated GFP signal in *atx-2(RNAi)* embryos compared to controls. It could be that the elevated levels of centrosomal ZYG-1 in *atx-2* mutants reflect a global increase in ZYG-1 throughout the cells. To test this possibility we compared overall ZYG-1 levels by measuring cytoplasmic GFP signals, but found no increase in the cytoplasmic levels of *atx-2(RNAi)* embryos. In fact, we noticed a small decrease (*p*=0.2) in the cytoplasmic GFP signal in *atx-2(RNAi)* embryos compared to controls, suggesting that increased centrosomal ZYG-1 levels are unlikely due to an increase in overall ZYG-1 expression. Similar results were also observed in *atx-2(ne4297)* mutants (**S6 Movie)**. Together, our data show that inhibiting ATX-2 results in elevated levels of ZYG-1 at centrosomes without affecting overall ZYG-1 levels. Thus, we speculate that elevated levels of centrosomal ZYG-1 in *atx-2* depleted embryos might partially compensate for the reduced activity of mutant ZYG-1 (P442L) in *zyg-1(it25)* (Kemp et al., 2007), restoring centrosome duplication in *zyg-1* mutants.

### ATX-2 negatively regulates centrosome size

Prior work has shown that SZY-20 negatively regulates centrosome size in a ZYG-1 dependent manner (Song et al., 2008). Here, our data indicate that ATX-2, acting downstream of SZY-20, affects centrosomal ZYG-1 levels. We thus tested if ATX-2 influences centrosome size as well. Quantitative IF using α-SPD-5 revealed that at first metaphase, *atx-2* mutant centrosomes possess twice as much SPD-5 as wild-type centrosomes (**Fig 5A, 5B**). Consistent with the results in *szy-20* mutants (Song et al., 2008), centrosomal SPD-2 levels are also significantly increased without affecting overall levels in *atx-2* mutants (**S5A and S5B Fig**). The centrosome factors SPD-5 and SPD-2, acting as a scaffold, are required for localization of other PCM factors including γ-Tubulin (Hamill et al., 2002; Kemp et al., 2004; Woodruff et al., 2015).

**Fig 5.**
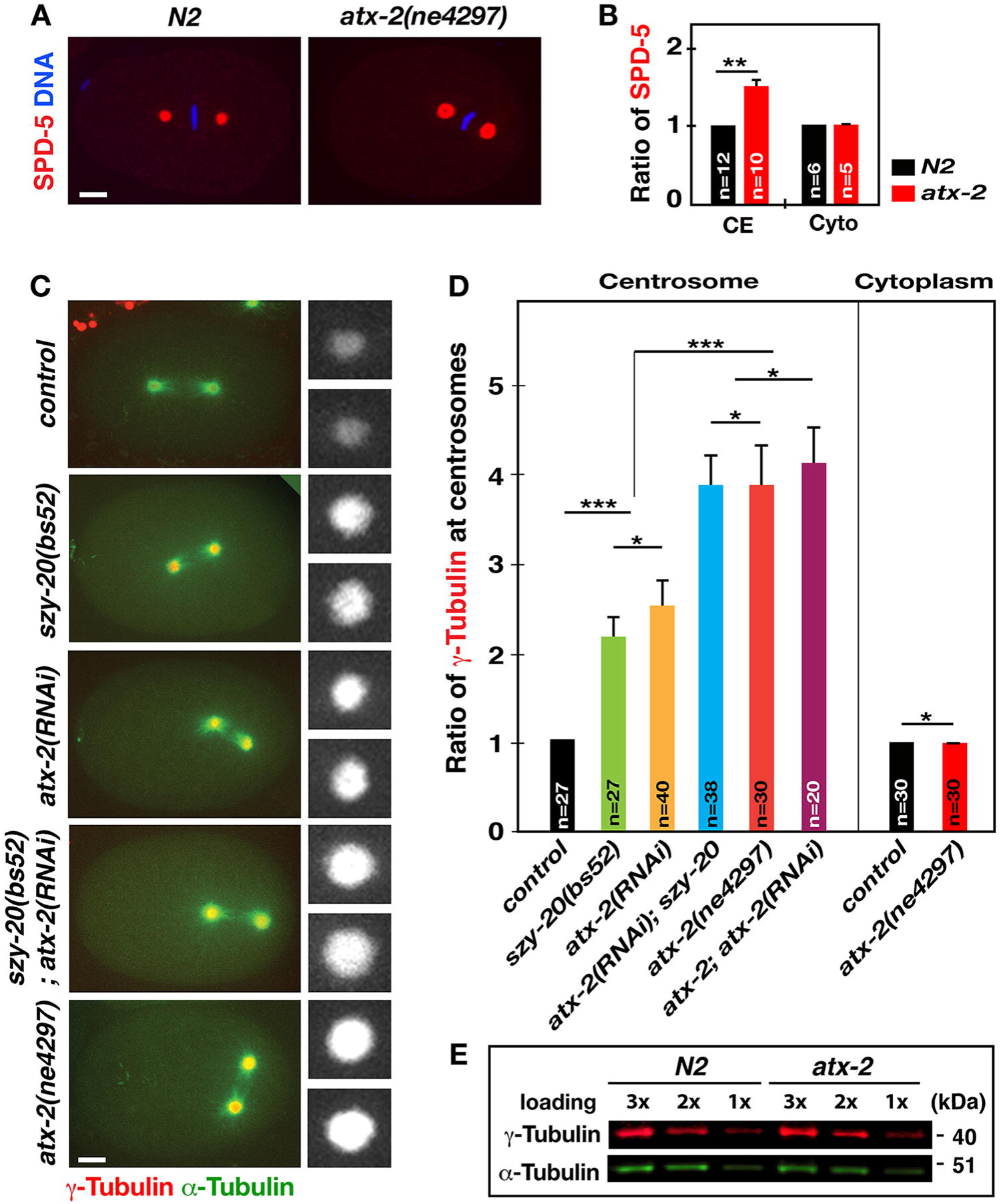
Knockdown of *atx-2* enhances levels of PCM factors at centrosomes. (A) *atx-2* embryos show increased centrosomal SPD-5 levels at the first metaphase. (B) Relative fluorescence intensity of centrosomal (CE) and cytoplasmic (Cyto) SPD-5 (\*\**p<*0.001). (C) Still images from time-lapse movies of embryos expressing mCherry-γ-Tubulin (Toya et al., 2010) and GFP-γ-Tubulin. Insets illustrate centrosomes magnified in 3-fold. (D) Measurements of fluorescence intensity of centrosomal and cytoplasmic γ-Tubulin at the first metaphase. Error bars are SD. (\*\*\**p*<0.0001; * *p*>0.005). Bar, 5 μm. (E) Quantitative immunoblot analysis using embryonic lysates reveals no significant changes in overall expression levels of γ-Tubulin (1.01 ± 0.1, n=10) in *atx-2* mutant embryos compared to *N2* control grown at 22°C. γ-Tubulin was used as loading control.

γ-Tubulin is a PCM factor that plays a key role in MT nucleation and anchoring (Hannak et al., 2002; O’Toole et al., 2012; Srayko et al., 2005). By 4D-time lapse confocal microscopy of embryos expressing mCherry-γ-Tubulin (Toya et al., 2010), we measured the fluorescent intensity of centrosomal mCherry signal at first metaphase (**Fig 5C, 5D**). Consistent with SPD-5 levels, *atx-2(RNAi)* embryos display a nearly two-fold increase in centrosomal γ-Tubulin, equivalent to the increase seen in *szy-20(bs52)* embryos (Song et al., 2008). A similar observation is made in a strain overexpressing GFP-γ-Tubulin (**S5C Fig**). Note that *szy-20(bs52)* embryos possess ~50% of normal ATX-2 levels, which is comparable with the 50% average knockdown of ATX-2 by feeding *atx-2(RNAi)* (**Fig 3E, 3K**).

Remarkably, *atx-2(ne4297)* mutant centrosomes show even greater increase (4-fold) than *szy-20(bs52)* or *atx-2(RNAi)* embryos, suggesting a dose-dependent correlation between the amount of ATX-2 and centrosomal γ-Tubulin. Compared to single knockdown by *szy-20(bs52)* or *atx-2(RNAi)*, co-depletion by *szy-20(bs52); atx-2(RNAi)* further exacerbated centrosomal enlargement, in a manner nearly equivalent to those in the strong loss-of-function mutation *atx-2(ne4297)* embryo. Thus, centrosomal γ-Tubulin levels correlate inversely with embryonic ATX-2 levels. Cytoplasmic mCherry-γ-Tubulin levels, however, are found to be similar in all embryos examined (**Fig 5D**). Furthermore, quantitative immunoblot confirms no significant effect on endogenous γ-Tubulin levels in *atx-2(ne4297)* compared to wild-type embryos (**Fig 5E)**. Therefore, our data show that centrosomal PCM levels (SPD-2, SPD-5, γ-Tubulin) correlate inversely with ATX-2 abundance, suggesting that ATX-2 negatively regulates centrosome size in proportion to its abundance, while not affecting overall cellular levels of centrosome factors.

### ATX-2 limits the number of microtubules emanating from the centrosome

The PCM factors, SPD-2, SPD-5 and γ-Tubulin, play a critical role in positively regulating the MT nucleating capacity of the centrosome (Hamill et al., 2002; Hannak et al., 2002; O’Toole et al., 2012; Srayko et al; 2005). As *atx-2* mutant embryos exhibit enlarged centrosomes with increased centrosomal SPD-5 and γ-Tubulin, we examined if *atx-2* mutant centrosomes affected MT nucleating capacity. To investigate MT nucleation, we used a strain expressing EBP-2-GFP to mark the plus-ends of growing MTs (Srayko et al., 2005) and acquired a series of 500 msec-interval snap shots at the center plane of first metaphase centrosomes (**Fig 6, S7 Movie**). First, we found that *atx-2* mutant embryos exhibit a three-fold increase (*p*<0.0001) in centrosomal EBP-2-GFP signal compared to wild-type controls (**Fig 6A, 6B**). To assess the level of MT nucleation by the centrosome, we measured the fluorescent intensity of EBP-2-GFP in line regions proximal (25 pixels) to the centrosome, finding that *atx-2* mutant centrosomes exhibit a two-fold increase (*p*<0.001) in MT nucleation (**Fig 6C**). Consistent with increased MT nucleation, kymograph analysis at the proximal region to the centrosome revealed a drastic increase (*p*=0.002) in the number of astral MTs emanating from *atx-2* mutant centrosomes over a 5-sec period, compared to controls (**Fig 6F**). Further, line scans of the kymograph in a 355° arc around the centrosome show that mutant centrosomes emanate increased numbers of spindle fibers as well as astral MTs (**Fig 6D**). Thus, *atx-2* mutant centrosomes possessing elevated levels of PCM factors nucleate the increased number of astral and spindle MTs.

**Fig 6.**
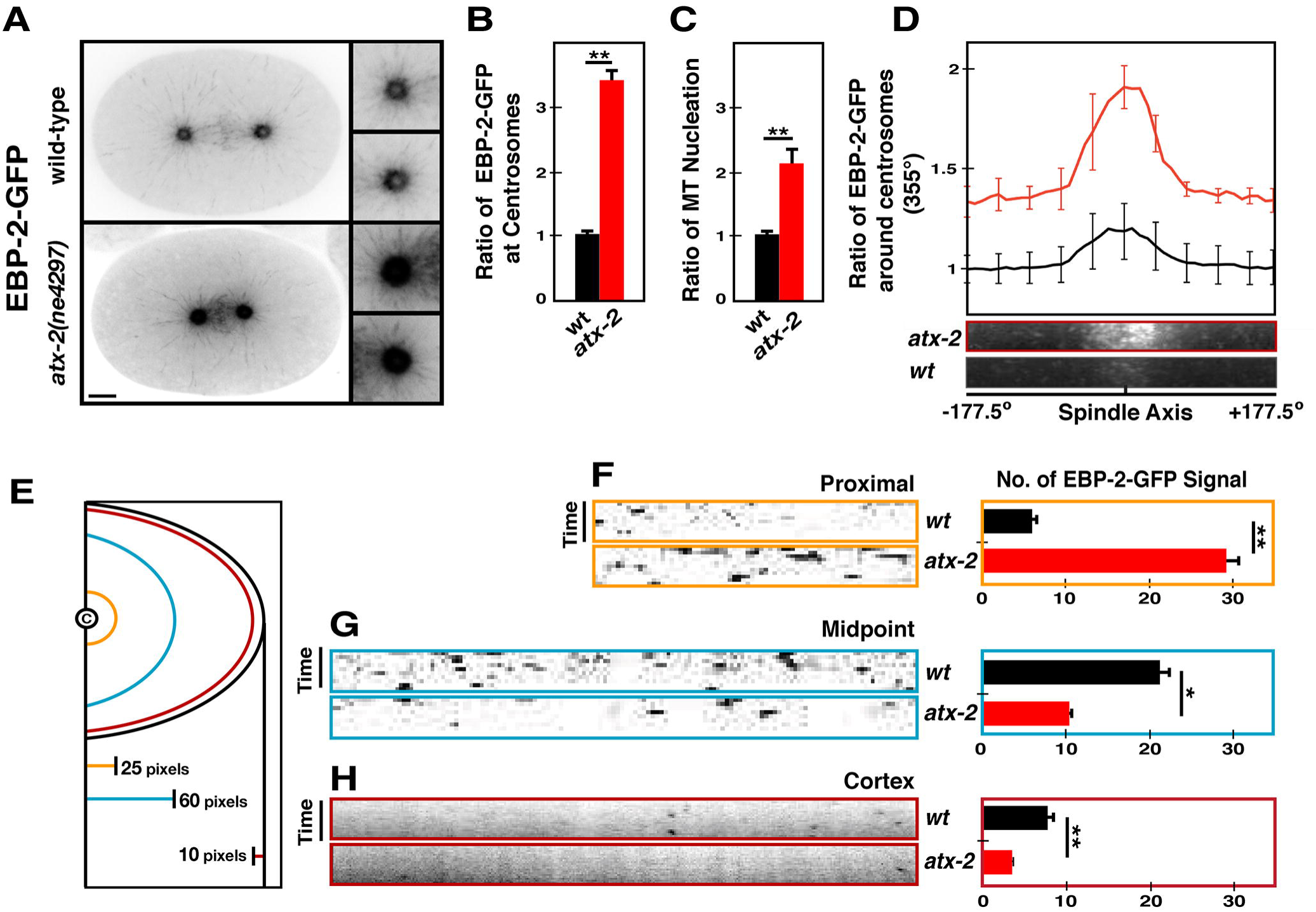
Loss of *atx-2* leads to increased MT nucleation and short MT formation. Time-lapse recordings of embryos expressing EBP-2-GFP reveal MT behavior. (A) 5-sec time projections of 500 msec interval time-lapse recording illustrate the tracks of MT growth over a 5-sec period. Note the short spindle and astral microtubules in *atx-2* embryos. Insets are magnified 3-fold. Bar, 5 μm. (B) Fluorescence intensity of centrosomal EBP-2-GFP at first metaphase centrosomes (n=18, \*\**p<*0.0001). (C) Relative levels of MT nucleation based on the fluorescent intensity of EBP-2-GFP in line regions (n=11, ** *p<*0.001; see Methods and Materials for quantification). (D) Line scans of the kymographs 355° arc (35 pixel radius) around centrosomes show that *atx-2* centrosomes emanate the significantly increased number of both spindle fibers and astral MTs. Note that spindle fibers appear wider and disorderly in *atx-2* mutants. (E) Schematic of kymograph analysis. Center circle illustrates the centrosome. Distance from the centrosome is shown in pixels (pixel = 0.151 μm) (F-H) Kymograph analysis of growing MTs at the (F) proximal [6.3 ± 0.7 (n=17, wt) vs 28.93 ± 1.2 (n=21, *atx-2*), \*\**p*=0.002] (G) midpoint [21.85 ± 1.33 (n=13, wt) vs 10.59 ± 0.54 (n=17, *atx-2*), \**p* = 0.029] from the centrosome, and near the (H) cortex [6.6 ± 0.12 (n=18, wt) vs 3.3 ± 0.15 (n=15, *atx-2*), \*\**p*=0.001] over a 5-sec period. Error bars are SD.

### Excessive MT nucleation leads to aberrant MT growth in *atx-2* mutants

Next we examined how these MTs nucleated by mutant centrosomes continue to grow out toward the cortex (**Fig 6E, 6G, 6H, S7 Movie**). To measure the number of MTs reaching the cortex, we used kymograph analysis over a 5-sec period and counted the number of EBP-2-GFP dots crossing an arc drawn 1.5 μm (10 pixels) inside the cortex. In mutant embryos, fewer MTs grew out to reach the cortex than in controls (*p*=0.001; **Fig 6H**). Consistently, only a small portion of MTs nucleated by mutant centrosomes grew beyond the midpoint between the centrosome and cortex compared to controls (*p*=0.029; **Fig 6G**). In fact, astral MT growth rates are significantly slower in mutant embryos (0.70 μm/sec ± 0.007) than in controls (0.88μm/sec ± 0.013, *p*=0.0014) (**S5E Fig**). Thus, MT growth appears to be impeded in the mutant embryo, presumably due to an excess of MT nucleation.

In *C. elegans* embryos, it has been shown that fast MT growth is subject to the MT stabilizing complex (ZYG-9/TAC-1) and the amount of free tubulin (Srayko et al., 2005). *atx-2* mutant embryos, however, exhibit a significant increase in both centrosomal and overall TAC-1 levels (**S5F and S5G Fig**), indicating that defective MT growth in *atx-2* mutants is unlikely due to insufficient MT stabilization. Studies in other systems also showed that the amount of free tubulin influences the rate of MT polymerization *in vitro* (Belmont et al., 1990; Walker et al., 1988) and such mechanism appears to be pertinent in *C. elegans* embryos (Lacroix et al., 2016; Song et al., 2008; Srayko et al., 2005). We thus hypothesized that *atx-2* mutant centrosomes possessing increased PCM nucleate more MTs, reducing the supply of free tubulin available for MT polymerization. Subsequently MT growth is interfered, resulting in shorter than normal MTs and cytokinesis failure. To test this, we generated *atx-2* mutants overexpressing GFP-Tubulin to see if increasing Tubulin levels in the *atx-2* mutant could partially rescue cytokinesis defects. As predicted, the incidence of cytokinesis failure is significantly (*p*<0.001) reduced in GFP-Tubulin overexpressing *atx-2* embryos compared to control mutant embryos (**Fig 7A**). This partial rescue of cytokinesis failure further led to a significant decrease (*p*<0.001) in embryonic lethality (**Fig 7B**), suggesting that MT growth in *atx-2* embryos is partly affected by limited supply of free tubulins, likely resulting from excessive MT nucleation. Next, we examined *atx-2* mutant embryos overexpressing GFP-γ-Tubulin, a PCM factor, to see if further enhancing MT nucleation by increasing γ-Tubulin levels could exacerbate cytokinesis defects through worsened MT growth. Consistent with our hypothesis, cytokinesis defects in mutant embryos were significantly increased by GFP-γ-Tubulin overexpression compared to control mutants, leading to an increase in embryonic lethality (**Fig 7A, 7B)**. Together, our data suggest that excessive MT nucleation and subsequent MT growth defects result in cytokinesis defects in *atx-2* mutants.

**Fig 7.**
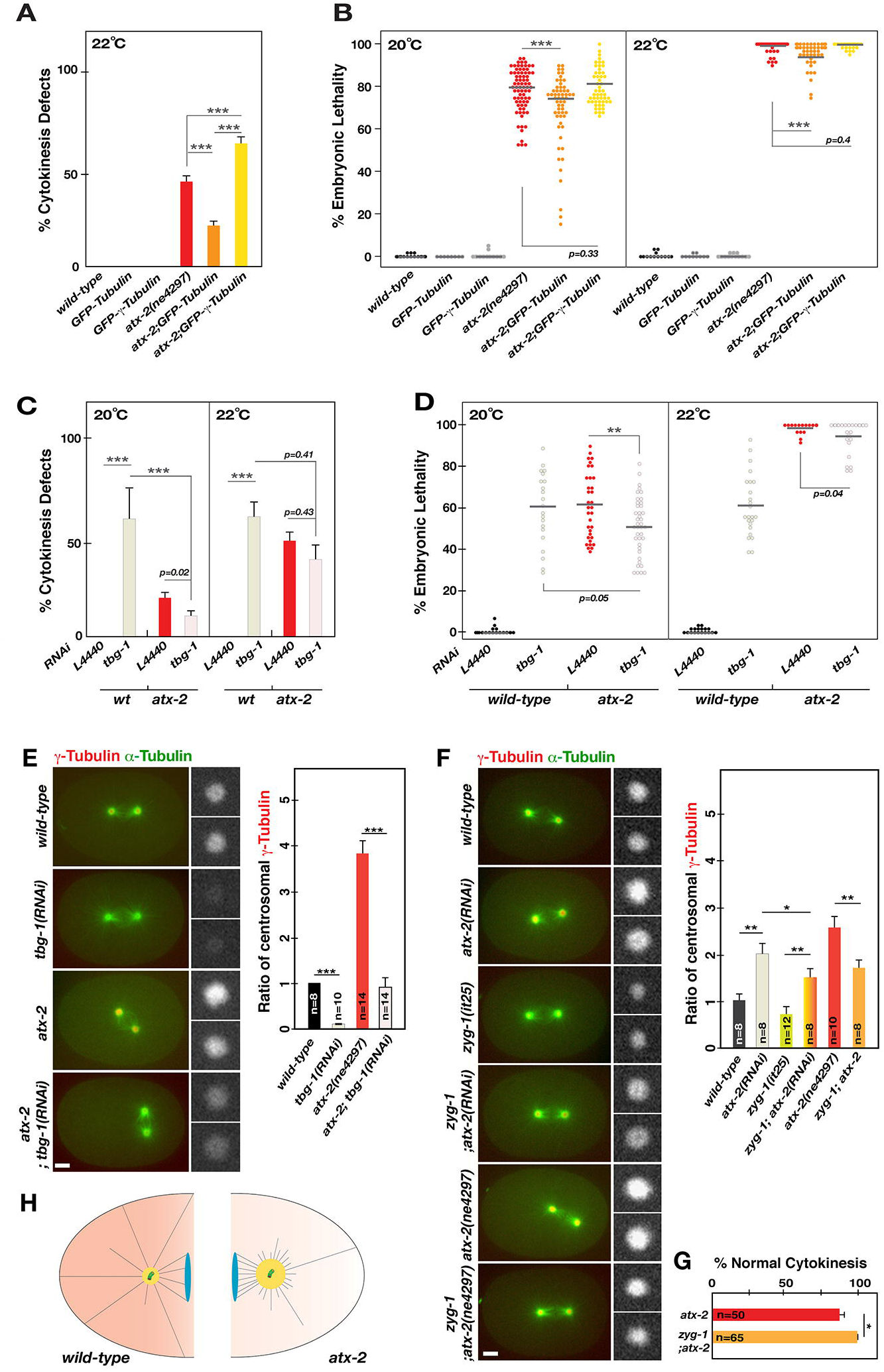
ATX-2 influences MT behavior through **γ-Tubulin.** (A) Cytokinesis events at first and second cell divisions were examined in wild-type or *atx-2* mutant embryos overexpressing GFP-Tubulin or GFP-γ-Tubulin at 22°C, the restrictive temperature for *atx-2* mutants. Compared to *atx-2* mutants (46%, n=490) alone, the percent of cytokinesis failure is significantly reduced in *atx-2* embryos overexpressing GFPγ-Tubulin (29%, n=280). In contrast, cytokinesis failure is enhanced in *atx-2* embryos overexpressing GFP-γ-Tubulin (68%, n=172), while wild-type embryos (n=63-109) overexpressing either GFP-protein exhibit no cytokinesis defects. (B) Embryonic lethality was scored at semi-restrictive (20°C) and restrictive (22°C) conditions for *atx-2* mutants. Overexpression of GFPγ-Tubulin in the mutant leads to a small but significant (\*\*\**p*< 0.0001) decrease in embryonic lethality: 78% (n=3852) and 98.6% (n=1949) in *atx-2* vs 69% (n=2776) and 95% (n=1829) in *atx-2;* GFPγ-Tubulin at 20 and 22°C, respectively. However, overexpressing GFP-γ-Tubulin leads to a small increase (*p*>0.05) in embryonic lethality: 81% (n=3087) and 99% (n=1149) at 20 and 22°C, respectively, compared to the *atx-2* mutant alone. A At 20°C, *tbg-1(RNAi)* in *atx-2* mutant embryos (10.4%, n=235) decreased cytokinesis defects compared to *atx-2* mutants (19.8%, n=270), while *tbg-1(RNAi)* produced cytokinesis failure (67%, n=331) in wild-type embryos. At 22°C, the same trends were observed: 38.8% (n=201) in *tbg-1(RNAi); atx-2,* 48.3% (n=179) in *atx-2*, 68% (n=380) in *tbg-1(RNAi)*. (D) At 20°C, *tbg-1(RNAi)* in the mutant leads to a small decrease (*p<*0.01) in embryonic lethality: 52% (n=2189) in *tbg-1(RNAi); atx-2,* 61.6% (n=2359) in *atx-2*, 60.6% (n=1935) in *tbg-1(RNAi)*. A similar observation was made at 22°C: 95% (n=975) in *tbg-1(RNAi); atx-2,* 98.7% (n=732) in *atx-2*, 63% (n=1456) in *tbg-1(RNAi)*. (A,C) Average values are shown, n= the number of embryos. (B,D) Horizontal bars indicate average values. Each dot represents a single animal, n= the number of progeny; \*\**p*<0.01; \*\*\**p*<0.001. (E) Embryos expressing GFP-γ-Tubulin and mCherry-γ-Tubulin are exposed to *tbg-1(RNAi)* at 22?C, the restrictive temperature for *atx*-2 mutants. While RNAi almost abolishes mCherry signal in wild-type, *tbg-1(RNAi)* reduces mCherry signal nearly to the wild-type level in *atx*-2 mutants. Relative levels of centrosomal - Tubulin are shown (\*\**p*<0.001). (F) Embryos grown at 20?C display centrosome-associated - Tubulin and microtubules at first metaphase, displaying that inhibiting ZYG-1 activity reduces centrosomal γ-Tubulin levels in *atx-2(RNAi)* or mutant embryos. (G) Quantification of centrosomal γ-Tubulin at the first metaphase. Relative levels of centrosomal γ-Tubulin are shown (* *p<* 0.05; \*\**p*<0.01; \*\*\**p*<0.001). (H) The *zyg-1(it25)* mutation restores normal cytokinesis to *atx-2* mutants at 20?C, the semi-restrictive condition for *atx-2(ne4297)*. (E, F) Insets illustrate centrosomes magnified in 3-fold. Error bars are SD. Bar, 5 μm. (H) In the wild-type embryo (left), SZY-20 acts upstream to promote levels of ATX-2, which negatively regulates centrosome size (ZYG-1, γ-Tubulin). A balance between positive and negative regulators establishes proper centrosome size and MT nucleating activity, contributing to normal MT growth. In *atx*-2 (or *szy*-20) mutant embryos (right), reduced levels of ATX-2 (red) disrupt this balance, thereby increasing centrosome size, which leads to increased numbers of MTs emanating from the centrosome and shortened MTs. Under normal conditions, ATX-2 limits centrosomal PCM levels to direct proper MT nucleating activity of the centrosome and MT growth.

### Cytokinesis defects in *atx-2* mutants result from excessive MT nucleation due to enlarged centrosomes

If defective cytokinesis results from elevated PCM levels at mutant centrosomes through excessive MT nucleation, reducing PCM levels should reconcile proper centrosomal PCM levels and MT nucleation, thereby restoring proper cytokinesis in *atx-2* mutants. Whereas *tbg-1(RNAi)* produces cytokinesis failure in wild-type embryos, reducing γ-Tubulin in *atx-2* embryos partially restores normal cytokinesis (*p<*0.001, **Fig 7C**), and embryonic viability (*p<*0.01, **Fig 7D**) to mutant embryos. Interestingly, not only did *tbg-1(RNAi)* rescue cytokinesis defects in *atx-2* mutants, but *atx-2* mutation also reduced cytokinesis failure in *tbg-1(RNAi)* embryos, suggesting a mutual suppression between *atx-2* and *tbg-1* (**Fig 7C, 7D)**: At 20°C, the semi-permissive temperature for *atx-2* mutants, single knockdown of *atx-2(ne4297)* and *tbg-1(RNAi)* produces 24% (n=91) and 66.5% (n=73), respectively, compared to 10.8% (n=57) of cytokinesis failure by double knockdown in *atx-2(ne4297)*; *tbg-1(RNAi)* embryos. Consistently, *atx-2(ne4297)*; *tbg-1(RNAi)* embryos exhibited centrosome-associated γ-Tubulin nearly to the normal level, although *tbg-1(RNAi)* almost abolished centrosomal γ-Tubulin in wild-type embryos (**Fig 7E)**. It appears that ATX-2 influences MT nucleation and MT dynamics through γ-Tubulin at the centrosome, suggesting an epistatic relationship between *atx*-2 and *tbg-1.* A prior study reported a mutual suppression between *szy-20* and *zyg-1* (Song et al., 2008). Given the genetic and functional relationship between *atx-2* and *szy-20*, we speculated that loss of *zyg-1* might have a similar effect on *atx-2.* At semi-restrictive conditions for the *atx-2* mutant, double homozygous mutants *zyg-1(it25); atx-2(ne4297)* produce a small but significant increase (*p*=0.02) in embryonic viability compared to the *atx-2(ne4297)* single mutant, accompanied by the restoration of normal cytokinesis (**Table 1**). Restoring normal cytokinesis appears to correlate with *zyg-1(it25)*-mediated restoration of centrosome size in *atx-2* embryos (**Fig 7F**). In fact, *zyg-1(it25)* embryos exhibit decreased levels of centrosomal γ-Tubulin at the permissive temperature 20°C where centrosome duplication occurs normally. Moreover, inhibiting ZYG-1 reduces centrosomal γ-Tubulin levels in *atx-2(RNAi)* or mutant embryos, which likely leads to the partial restoration of normal cytokinesis and embryonic viability in ATX-2 depleted embryos (**Fig 7F, 7G, Table 1**).

Together, our results suggest a model in which loss of ATX-2 leads to elevated PCM levels at centrosomes and excessive MT nucleation, resulting in MT growth defects and subsequent cytokinesis failure (**Fig 7H**). Consistently, embryos depleted of ATX-2 exhibit a prominent cytokinesis failure phenotype, accompanied by enlarged centrosomes that cause excessive MT nucleation and aberrant MT growth, which results in cytokinesis failure and embryonic lethality. Thus, the proper level of centrosomal PCM factors is critical for normal MT nucleating activity to support normal MT dynamics. In this pathway, ATX-2 acts upstream to establish the proper centrosome size and MT nucleating activity.

## Discussion

### ATX-2 forms a complex with SZY-20 and acts as a negative regulator of centrosome assembly in *C. elegans* embryos

In this study, we identified that ATX-2, together with PAB-1, physically associates with SZY-20 *in vivo*. While RNA-binding proteins ATX-2 and SZY-20, each containing unique RNA-binding motifs (LSm, SUZ and SUZ-C, respectively), form a complex, these two proteins appear to physically interact independently of RNA. While it remains unknown what RNA molecules associate with these RNA-binding proteins, each RNA-binding protein in a ribonucleoprotein (RNP) complex might recruit a specific group of RNA through each own RNA-binding motif. In fact, Ataxin-2 has been shown to bind directly to mRNAs through its LSm domain and promote the stability of transcripts independently of its binding partner, poly (A)-binding protein (PABP) in flies and humans (Satterfield and Pallanck 2006; Yokoshi et al., 2014). Our prior study in *C. elegans* embryos showed that SUZ and SUZ-C RNA-binding motifs in SZY-20 exhibit RNA-binding capacity *in vitro* and that mutating these domains perturbs *in vitro* RNA-binding of SZY-20 and its capacity to regulate centrosome size *in vivo* (Song et al., 2008). However, there is no evidence that RNA-binding role of ATX-2 is directly involved in centrosome regulation, while *C. elegans* ATX-2 is shown to function in translational regulation during germline development (Ciosk *et al*., 2004), In a multi-protein complex, ATX-2 and SZY-20 function closely to regulate cell division and centrosome assembly. While knocking down ATX-2 phenocopies a loss of function *szy-20(bs52)* mutation, the different degree of phenotypic penetrance in *atx-2(RNAi),* a strong loss of function *atx-2(ne4297)* mutation, and a hypomorphic *szy-20(bs52)* mutation suggests a dose-dependent regulation of ATX-2. Double knockdown by combining *atx-2(RNAi)* and *szy-20(bs52)* mutation further enhances embryonic lethality, cytokinesis failure, the restoration of centrosome duplication to *zyg-1(it25)* embryos and the levels of centrosome-associated factors (ZYG-1, SPD-5, γ-Tubulin). Furthermore, the *atx-2(ne4297)* mutation produces a similar effect to that of *atx-2(RNAi)* combined with the *szy-20(bs52)* mutation. Our data thus suggest that *atx-2* exhibits a positive genetic interaction with *szy-20* in regulating cell cycle and centrosome assembly. In this pathway, SZY-20 acts upstream of ATX-2 to promote ATX-2 levels, with both acting upstream of *zyg-1.* We propose that SZY-20 influences centrosome size and MT dynamics indirectly through ATX-2 (Song et al., 2008). While it remains unclear whether *C. elegans* ATX-2 directly acts on ZYG-1, it has been shown that Plx4 (a *Xenopus* homolog of Plk4/ZYG-1) forms a complex with Atxn-2 (*Xenopus* homolog of ATX-2; Hatch et al., 2010). Although a direct role for Atxn-2 in centrosome assembly has not been demonstrated, a physical connection between *Xenopus* Atxn-2 and Plx4 suggests a possible role of *Xenopus* Atxn-2 in centrosome assembly, via a mechanism that seems likely to be conserved between nematodes and vertebrates.

### ATX-2 negatively regulates centrosome size in a dose-dependent manner

Our data indicate that ATX-2 acts as a negative regulator of centrosome size. A priori, increased PCM levels at *atx-2* mutant centrosomes could be achieved by several mechanisms. First, centrosome factors might be overexpressed in *atx-2* mutant cytoplasm via translational control, leading to increased recruitment of these factors to the centrosome by equilibrium, as shown by Decker et al (2011). Second, *atx-2* mutants might enhance the recruitment of factors to centrosomes post-translationally, without affecting overall levels of these factors. Third, loss of ATX-2 might promote local translation near centrosomes, leading to locally enriched centrosome factors. The first scenario is unlikely because our quantitative analyses reveal no significant changes in overall levels of centrosome factors (SPD-2, SAS-6, γ-Tubulin) or cytoplasmic levels of ZYG-1 by loss of ATX-2. Our current data do not differentiate between the second and third scenarios, but certain observations in *C. elegans* and other systems are consistent with the latter. It has been shown that Ataxin-2 assembles with polyribosomes and that ribosomes are associated with the MT cytoskeleton (Hamill et al., 1994; Satterfield and Pallanck 2006). Furthermore, neuronal RNA granules are shown to contain translational machinery, allowing local translation upon arrival of transcripts at the right location (Shiina et al., 2005). We also observed ribosomal protein S6 spatially associated with *C. elegans* centrosomes (**S3D Fig**), suggesting the possibility of local translation around the centrosome. Indeed mRNA localization linked to the cytoskeleton and the following local translation is shown to be an efficient means of concentrating proteins at the functional site (Groisman et al., 2000; Huttelmaier et al., 2005; Kindler et al., 2005). For example, spatial control of β-actin translation is executed by localizing its transcripts to actin-rich protrusions, which facilitates neuronal outgrowth (Huttelmaier et al., 2005). Also, RNA-binding protein (CPEB/maskin) mediated localization of cyclin B1 transcripts to the mitotic apparatus leads to locally enriched Cyclin B proteins in the vicinity of spindles and centrosomes, supporting cell cycle progression (Griosman et al., 2000). Together such mechanism has been demonstrated to provide a tight control for temporal and spatial translation. How then, might ATX-2 regulate translation? For negative translational control, it has been proposed that Ataxin-2 binding to PABP inhibits translation by blocking the interaction between PABP and translational machinery or by directly blocking translation of mRNA targets at the initiation stage (Besse and Ephrussi, 2008; Roy et al., 2004; Satterfield and Pallanck 2006; Yokoshi et al., 2014). Alternatively, ATX-2 may facilitate the interaction between a microRNA and its mRNA target, leading to translational inhibition (McCann et al., 2011). ATX-2 might also mediate translational control via its RNA-binding role as a component of the RNP complex comprising SZY-20/ATX-2/PAB-1. In fact, *C. elegans* PAB-1 is shown to associate with stress granules and processing bodies (P-bodies) that have been implicated in translational repression (Gallo et al., 2008). It has been observed that RNP complexes including P-bodies are involved in subcellular targeting of RNAs and precise timing of local translation at the final subcellular destination (Barbee et al., 2006; Besse and Ephrussi, 2008). Further identification of specific RNAs that bind ATX-2 and/or SZY-20 will help to understand how RNA-binding role of an ATX-2 associated RNP complex plays a role in defining proper centrosome size.

### ATX-2 contributes to proper MT nucleating activity of centrosomes and MT growth through γ-Tubulin

We have shown here that ATX-2 plays an essential role in cell division. *atx-2* mutant embryos with enlarged centrosomes exhibit multiple cell division defects including cytokinesis failure and aberrant spindle positioning. In animal cells, contact between astral MT and the cortex is critical for the initiation of the cleavage furrow that is required for proper cytokinesis (Green et al., 2012; Normand and King, 2010). Thus cell cycle defects in *atx-2(ne4297)* embryos might associate with aberrant MT behavior, likely due to enlarged centrosomes. Recent work showed an additional role of ZYG-1 in regulating centrosome size, independently of its role in centriole duplication (Song et al., 2008). PCM factors SPD-2, SPD-5 and γ-Tubulin are known to positively regulate the MT nucleating activity of the centrosome (Hamill et al., 2002; Hannak et al., 2002; O’Toole et al., 2012; Wueseke et al., 2015). *atx-2* mutant centrosomes possessing increased (2-5 folds) levels of centrosome factors nucleate the drastically increased number of MTs. Such increase in MT nucleation by mutant centrosomes may lead to a substantial reduction in cytoplasmic free tubulins available for timely MT polymerization. Given the rapid cell divisions (~20 min/cell cycle) and high MT growth rate (0.88 µm/sec) in early *C. elegans* embryos, timely and sufficient supply of free tubulins in the cytoplasm must be immensely critical for proper cell cycle progression. In support of this, either overexpressing Tubulin or reducing positive regulators (ZYG-1 or γ-Tubulin) of MT nucleation partially restores normal cytokinesis, MT growth and embryonic viability to *atx-2* mutants. Partial restoration of normal MT-dependent processes correlates with proper levels of centrosomal γ-Tubulin. Therefore, enlarged centrosomes are likely to be a primary cause of embryonic lethality in *atx-2* mutants. Mutant centrosomes possess significantly increased levels of γ-Tubulin, and reducing its levels at mutant centrosomes partially reinstalls free tubulins available for MT growth, which in turn restores normal cell divisions and embryonic viability in *atx-2* mutants. Our data suggest that ATX-2 contributes to MT nucleation and MT dynamics, in part, through γ-Tubulin at centrosomes, although it remains unknown how RNA-binding protein, ATX-2, regulates centrosomal levels of γ-Tubulin. Given our observation that overall levels of γ-Tubulin are not altered in *atx-2* mutant embryos, it is curious how γ-Tubulin levels are locally enriched at the centrosome, perhaps via the regulation of recruitment to the centrosome or locally enriched translational control. Thus, it will be interesting to see if *tbg-1* transcripts are elevated at the proximity of centrosomes by *in situ* hybridization. A recent study in *Xenopus* egg extracts reported that RNase treatment leads to defective spindle assembly due to hyperactive MT destabilizer MCAK (the ortholog of *C. elegans* KLP-7) by depleting RNA, suggesting a direct involvement of RNA in regulating MT organization (Jambhekar et al., 2014). Interestingly, we also found that *atx-2* mutants show increased levels of KLP-7 at centrosomes and cytoplasm (**S5H Fig**), and that *atx-2* and *klp-7* exhibit a synergistic effect on MT dynamics and embryonic lethality (**S6 Fig**).

Another centriole factor, SAS-4, positively regulates centrosome size in *C. elegans* (Kirkham et al., 2003). Interestingly, while we identify ATX-2 as a negative regulator of centrosome size, depleting ATX-2 had no effect on centriolar SAS-4 levels (**S4G Fig**), suggesting that ATX-2 regulates centrosome size independently of SAS-4, perhaps through a separate pathway. Thus, a balance between positive (e.g., SAS-4) and negative (e.g., ATX-2) regulators may contribute to establish proper centrosome size and MT nucleating activity, which in turn influences MT dynamics during the cell cycle. Improper levels of ATX-2 disrupt this balance, resulting in deregulated MT dynamics and subsequently abnormal cell divisions.

In summary, our work uncovers a role for ATX-2, the *C. elegans* ortholog of human Ataxin-2 in regulating centrosome size and MT cytoskeleton. In this pathway, SZY-20 positively regulates levels of ATX-2, which contributes to defining the proper centrosome size, leading to proper levels of MT nucleation and subsequent MT growth. While human Ataxin-2 and its poly Q stretch have been implicated in spinocerebellar ataxia (Lastres-Becker et al., 2008), it remains largely unknown how this RNA-binding protein is linked to human diseases. Many RNA-binding proteins in neuronal RNA granules have been implicated in neuronal functions and growth via controlled local translation (Bette and Ephrussi, 2008; Shiina et al., 2005), suggesting a link between RNA-binding role and neuronal activity. In this regard, our work provides insights into a mechanistic link between the RNA-binding role of Ataxin-2 and the MT cytoskeleton.

## Materials and Methods

### Strains and Genetics

All *C. elegans* strains were grown on MYOB plates seeded with *E. coli* OP50, and were derived from the wild-type Bristol N2 strain (Church et al., 1995). All strains were maintained at 18 or 20°C unless otherwise indicated. Transgenic strains were generated by standard particle bombardment transformation (Praitis et al., 2001) and the CRISPR/Cas-9 method (**S2 Table**, Dickinson et al., 2015; Paix et al., 2015). The list of worm strains used in this study is shown in **S1 Table**. Standard genetics were used for strain construction and genetic manipulations (Brenner, 1974). RNAi feeding was performed as described and L4440 clone with the empty dsRNA-feeding vector served as a negative control (Kamath et al., 2003).

### Cytological Analysis

For immunostaining, we used the following antibodies at a 1:400-3000 dilution: α-ATX-2 (Ciosk et al., 2004), DM1A (Sigma), α-GFP (Roche), α-SAS-4 (Song et al., 2008), α-SPD-2 (Kemp et al., 2004), α-SPD-5 (Hamill et al., 2002), α-SZY-20 (α-S20N: Song et al., 2008), and secondary antibodies Alexa Fluor 488 and 568 (Invitrogen). Affinity-purified rabbit polyclonal antibodies were generated (YenZym) against the following peptides: for γ-Tubulin (**S4D Fig,** aa31-44): Ac-HGINERGQTTHEDD-amide, for ZYG-1 (**S4E Fig,** aa671-689): Ac-PAIRPDDQRFMRTDRVPDR-amide.

Immunofluorescence and confocal microscopy were performed as described (Song et al., 2008). For confocal microscopy, MetaMorph software (Molecular Devices) was used to acquire images from a Nikon Eclipse Ti-U microscope equipped with a Plan Apo 60 X 1.4 NA lens, a Spinning Disk Confocal (CSU X1) and a Photometrics Evolve 512 camera. Fluorescence intensity measurements were made using the MetaMorph software, and image processing with Photoshop CS6.

For quantification of centrosomal signals, the average intensity within a 25-pixel (pixel = .151 μm) diameter region was measured throughout a region centered on the centrosome. For each region, a focal plane with the highest average intensity was recorded. The same sized regions were drawn in the cytoplasm for cytoplasmic signal, and outside the embryo for background subtraction unless otherwise indicated. For centriolar signals, analysis was done the same way, but a 5-pixel diameter region was used.

For EBP-2-GFP movies, pre-mitotic embryos were selected and monitored under DIC until before metaphase I. Upon entrance into metaphase, 61 images were captured every 500 msec for a 30 sec period. For spindle MTs, a 355° circular region was drawn around each centrosome, and kymographs were generated for the first 5 seconds of recordings. Linescan measurements of the kymographs were taken to obtain average EBP-2 signal along every point of the circular region. Values for each linescan were averaged and plotted on a graph. To measure MT nucleation, the fluorescence intensity was quantified at a line region drawn in a semi-circle (25 pixel radius) around the centrosome. Linescan (5 pixel wide) measurements were taken in 5-sec time projections from recordings. Pixel intensities were generated for each point along the line and averaged for comparison. For background subtraction, cytoplasmic intensity in the same size region was used. To quantify the number of growing astral microtubules proximal to the centrosome, line regions were drawn in a semi-circle (25 pixel radius) around the centrosome. For midpoint measurements, line regions were drawn at a radius of 60 pixels from the centrosome. Kymographs were generated for the first 5 seconds of each movie. Individual pixel measurements from the kymographs were obtained, and pixels with GFP intensity over 40 were counted. Cortex measurements were obtained by generating 5 sec kymographs of a region drawn at 1.5 μm inside of the cell cortex. The number of EBP-2 signal was counted manually. For MT polymerization rate, 4.5 μm-long lines were drawn along the individual EBP-2-GFP tracks outward to the cortex. Kymographs were generated, and rate was calculated by distance over time (μm/sec).

### Immunoprecipitation and Quantitative Western Analysis

20-25 μl of Dynabeads Sheep-anti-Rabbit, Protein A magnetic beads (Invitrogen) or mouse-anti-GFP magnetic beads (MBL) were used per IP reaction. Beads were washed 2x 15 min in 1 ml PBS with 0.1% Triton-X (PBST). For SZY-20 IP, beads were incubated overnight at 4°C in a ratio of 1mg/ml (α-SZY-20 or α-HA/beads volume) and 3 mg/ml (α-IgG/beads volume). Embryos were collected by bleaching worms grown in liquid culture, snap-frozen in liquid nitrogen, and stored at -80°C until use. Embryo pellets were ground up in microcentrifuge tubes in equal volumes of worm lysis buffer (50 mM HEPES; pH 7.4, 1 mM EDTA, 1 mM MgCl_2_, 200 mM KCl, 10% glycerol; Cheeseman et al., 2004) containing cOmplete protease inhibitor cocktail (Roche) and MG132 (Tocris). For RNase A or RNase Inhibitor treatment, RNase (10 μg/ml, Roche) or RNase Inhibitor (200U/ml, Roche) was added to lysis buffer prior to grinding. Embryos were ground for 5 min, and sonicated for 3 min. The lysate was then centrifuged at 4°C for 2x 20 min at 15K rpm in a desktop centrifuge, and the supernatant were collected. For enzyme reaction, lysates were incubated for 15 min at room temperature (RT). Protein concentration was determined and adjusted before IP. Beads were washed for 2x 15 min with PBST + 0.5% BSA at RT, followed by 1x 15 min wash with 1X worm lysis buffer. Beads were resuspended in 1X lysis buffer, mixed with embryonic lysates and incubated at 4°C for 1 hour. Following IP, beads were washed 2x 5 min in PBST. Samples were dissolved in 2X Laemmli Sample Buffer (Sigma) with 10% β-mercaptoethanol, fractionated on a 4-12% NuPAGE Bis-Tris Gel (Invitrogen) and blotted to nitrocellulose membrane using the iBlot Gel Transfer System (Invitrogen). α-ATX-2 (Ciosk et al., 2004), α-GFP (Roche), mouse α-HA (Sigma), α-SAS-6 (Song et al., 2011), α-SPD-2 (Song et al., 2008), α-SZY-20 (Song et al., 2008), DM1A (Sigma), α-TAC-1 (Hadwiger et al., 2010) and α-γ-Tubulin at 1:400-2500 dilution, and IRDye secondary antibodies (LI-COR Biosciences) were used at 1:10,000 dilution. Blots were imaged on a Licor Odyssey IR scanner, and analyzed using the Odyssey Infrared Imaging System (LI-COR Biosciences). Mass spectrometry analysis was performed as described (Song et al., 2011).

## Acknowledgements

We are grateful for the assistance of Mass Spectrometry Facility at National Institutes of Health. We also thank Song lab members for experimental assistance, David Weisblat for helpful discussion, Craig Mello, Kevin O'Connell, Ahna Skop, Masaki Shirayama and *Caenorhabditis* Genetics Center for worm strains and RNAi stains.

## Supporting information

**S1 Fig. ATX-2 and PAB-1 physically interacts with SZY-20.** (A) The amino acid sequences of peptides identified by MS/MS analysis and matched to those of ATX-2 and PAB-1 are highlighted in red. In comparison, MS analyses of the control IP showed only 2% peptide coverage for ATX-2 and 2.3% for PAB-1. The representative MS/MS spectra of two selected tryptic peptides (indicated in grey box) for each protein are shown below. In ATX-2, ***Q\**** in blue indicates the position that is mutated to stop codon in the *atx-2(ne4297)* strain (CAG to TAG), resulting in 128 aa deletion at the C-terminus. (B) ATX-2 is undetectable in pull-down from *szy-20(bs52)* lysates, while ATX-2 co-precipitates with SZY-20 in both RNase inhibitor or RNaseA treated wild-type lysates. Total RNAs were extracted from lysates treated with either RNase Inhibitor or RNase A using TRIzol (Thermo Fisher), analyzed by agarose gel electrophoresis: RNase A treatment efficiently degraded RNAs in lysates, whereas lysates treated with RNase Inhibitor exhibit bands of 18S and 28S ribosomal RNAs. In pull-down, the truncated form (dSZY-20 >) of SZY-20 in *szy-20(bs52)* mutants is possibly masked due to heavy detection of IgG heavy chain (<) near 51kDa. (C) To avoid heavy IgG detection, we performed α-GFP based IP assay using SZY-20-GFP embryonic lysates mixture with N2, *szy-20(bs52), atx-2(ne4297)* lysates. The truncated form of SZY-20 (dSZY-20) is undetectable in pull-down from SZY-20-GFP embryonic lysates, while the full-length SZY-20 co-precipitates with SZY-20-GFP, suggesting that the C-terminus of SZY-20 is required for the complex formation. (D) In *atx-2(ne4297),* the C-terminal truncation of ATX-2 (dATX-2) alleviates its interaction with SZY-20. dATX-2 signal is dramatically diminished in SZY-20 co-precipitates after 1x wash (1x w), compared to no wash (0x w) following IP, suggesting that the C-terminus of ATX-2 influences physical interaction with SZY-20. (E) Pull-down assays with α-GFP, using embryonic lysates from a strain expressing GFP fused to the C-terminus of SZY-20, PAB-1, ATX-2 and RPN-12. N2, and RPN-12-GFP are used as negative control. Consistent with the result in *szy-20(bs52)* embryonic lysates (**Fig 1**), ATX-2 is undetectable in pull-down from SZY-20-GFP lysates, further supporting that the C-terminus of SZY-20 is important for physical association with ATX-2. Both ATX-2 and SZY-20 are detected in pull-down from PAB-1-GFP and ATX-2-GFP lysates. In contrast, neither ATX-2 nor SZY-20 is detected in N2 and RPN-12-GFP lysates. For input, 5-10% of the total embryonic lysates were loaded for comparison.

**S2 Fig. Subcellular localization of ATX-2, GFP-PAB-1 and SZY-20.** (A) *pab-1(RNAi)* results in abnormal spindle positioning. (B) Wild-type embryos co-labeled for ATX-2 and SZY-20 illustrate that only a small fraction (~13%, n=11) of ATX-2 and SZY-20 foci coincide in the cytoplasm in early embryos (top). The embryo in Fig 3C is shown in each channel and the overlay (bottom). Arrows indicate colocalization. (C) Embryos co-labeled for ATX-2 and GFP-PAB-1 illustrate that cytoplasmic foci of ATX-2 and PAB-1 partially coincide (~20%, n=13) (top). The embryo in Fig 3D is shown in each channel and the overlay (bottom). Arrows indicate co-localizing cytoplasmic foci. (D) GFP-PAB-1 expressing embryos co-stained for GFP and SZY-20 illustrate that GFP-PAB-1 and SZY-20 partially colocalize (~3%, n=3) in the cytoplasm, and that SZY-20 expression is unaffected by *atx-2* or *pab-1(RNAi)*. Colocalization was quantified using the MetaMorph software. Circular regions with a diameter of 60 pixels were drawn at different positions within the cytoplasm. Threshold intensity values were set for each wavelength, and the percentage of overlapping pixels from each wavelength above the defined threshold values was calculated. Boxes for the overlays are magnified (2-2.5x) views. Bars, 5 μm.

**S3 Fig. *atx-2* mutation restores centrosome duplication in *zyg-1(it25)* embryos.** (A-B) *atx-2(ne4297)* mutation restores bipolar spindle formation to *zyg-1(it25)* embryos. At the second cell cycle, *zyg-1(it25); atx-2(ne4297)* double homozygous mutant embryos exhibits a cytokinesis failure with tetra-poles, indicating successful centrosome duplication. (B) Quantification of centrosome duplication based on the bipolar spindle formation at 22 and 23°C, semi-restrictive conditions for *zyg-1(it25)*. (C) Depleting CAR-1, an RNA-binding protein, does not restore centrosome duplication to *zyg-1(it25)* mutants. It is shown that CAR-1, another known RNA-binding protein, is required for normal cytokinesis (Audhya et al., 2005; Squirrell et al 2006). At 24°C, *car-1(RNAi)* in *zyg-1(it25)* results in 100% of monopolar spindles at the second mitosis, while *atx-2(RNAi)* restores > 80% of centrosome duplication to *zyg-1(it25)*. Thus, RNA-binding proteins ATX-2 and SZY-20 play a specific role in negatively regulating centrosome duplication. B Ribosomal subunit S6 partially coincides with γ-Tubulin at centrosomes in wild-type embryos. Insets are magnified 5-fold. Bar, 5 μm.

**S4 Fig. Localization of centrosome factors.** (A-C) The strain expressing an N-terminal HA tagged ZYG-1 at endogenous levels from the native genomic locus is generated using the CRISPR/Cas-9 method (**S2 Table** for Methods): γ-Tubulin as centrosome marker. HA-ZYG-1 localizes to centrioles, coinciding with anti-ZYG-1 labeling (A) in a cell cycle dependent manner (B): undetectable at first (a) and second (c) metaphase, and highest at late mitosis (b, c, d). (C) *zyg-1(RNAi)* almost abolishes HA-ZYG-1 signals. Compared to controls, *atx-2(RNAi)* embryos exhibit increased levels (~2-fold, n=10) of centrosome-associated HA-ZYG-1 signal at late mitosis. (D) The specificity of γ-Tubulin antibody: Immunostaining and immunoblot reveal that anti-γ-Tubulin specifically detects endogenous γ-Tubulin, as *tbg-1(RNAi)* leads to a significant reduction in γ-Tubulin signals. (E) The specificity of ZYG-1 antibody: ZYG-1 is enriched at centrioles. *zyg-1(RNAi)* results in a drastic reduction in both centriolar and cytoplasmic ZYG-1 signals. *zyg-1(RNAi)* leads to monopolar mitotic spindles at the second mitosis (bottom right). (F) SPD-5 is not required for ZYG-1 localization at centrosomes. ZYG-1 localizes to centrosomes in *spd-5(RNAi)* embryos that exhibit cell cycle arrest at the first mitosis (Hamill et al., 2002). Quantification of ZYG-1 levels at centrosomes reveals no significant change in ZYG-1 levels by loss of *spd-5*: Mitotic arrest in *spd-5(RNAi)* embryos is likely to contribute to more frequent detection of higher fluorescence intensity as ZYG-1 levels at centrosomes peak at late mitosis. (G) SAS-4 levels are not affected by loss of *atx-2*: SAS-4 stained embryos display SAS-4 localization at early cell cycle stages. Note cell cycle defects (e.g., DNA segregation, spindle positioning, cytokinesis) in *atx-2* mutants. Quantification of SAS-4 signals shows that *atx*-2 mutant embryos exhibit similar levels of SAS-4 at centrosomes (CE) and cytoplasm (Cyto) to those of wild-type embryos at metaphase. Insets are magnified 3-fold. Bar, 5 μm.

**S5 Fig**. **Loss of ATX-2 influences localization and levels of centrosome factors.** Centrosomal SPD-2 levels (A) are significantly increased without affecting overall levels (B) in *atx-2* embryos. (C) *atx-2(RNAi)* embryos display a nearly 3-fold increase in centrosomal GFP-γ-Tubulin but no change in cytoplasmic levels. (D) Line scans of the kymographs on embryos immunostained for MTs show that *atx-2* mutant embryos exhibit fewer MT-signals at the midpoint (9.6 μm away from centrosomes) and near the cortex (1.5 μm inside cortex) compared to wild-type controls, which is consistent with the analyses performed using EBP-2-GFP in Fig A) Each dot on the graph represents an embryo. Horizontal bars indicate average values, \*\*\**p*<0.001. (E) Polymerization velocity: Kymographs created along the growing MT. Lines (4.5 μm) were drawn along the individual EBP-2-GFP track extending outward to the cortex. Astral MT growth rates are significantly lower in mutant embryos (0.70 μm/sec ± 0.007) than in controls (0.88 μm/sec ± 0.013, \*\**p*=0.0014). (F) *atx-2* mutant embryos exhibit increased levels of TAC-1. Quantitative immunoblot analysis using embryonic extracts of wild-type and *atx*- 2*(ne4297)* mutants expressing GFP-TAC-1 (Le Bot et al., 2003) with -GFP and -TAC-1 (Hadwiger et al., 2010) shows that mutant embryos possess increased levels of both GFP-TAC-1 and endogenous TAC-1. Additional analysis using wild-type and *atx-2* embryos confirms that *atx-2* embryos contain elevated levels of TAC-1 (n=5). < indicating a truncated form of ATX-2 (dATX-2) in *atx-2* mutants. (A, F) Tubulin as loading control. (G) Z-projections of embryos expressing GFP-TAC-1 at first metaphase. The *atx*-2 mutant embryo displays a 3-fold increase (\*\**p*<0.001) in centrosomal GFP-TAC-1 levels. (H) Z-projections of embryos expressing KLP-7-GFP at first metaphase. In *atx-2(RNAi)* embryos, KLP-7-GFP signal at centrosomes (122.75 ± 27.72, n=8) is increased by ~2-fold compared to controls (71.32 ± 13.37, n=10, \*\**p*< 0.001). Kinetochore-associated KLP-7 reveals misaligned chromosomes (arrows) in *atx-2(RNAi)* embryos. (G, H) Centrosome (CE) and cytoplasm (Cyto); Insets are magnified 3-fold. Error bars are SD. Bar, 5 μm.

**S6 Fig**. **Loss of ATX-2 leads to defective MT growth likely due to excessive MT nucleation.** (A) 5-sec time projections of 500 msec interval live imaging of embryos expressing EBP-2-GFP at first metaphase illustrate the effects on MT behavior in RNAi-treated *atx-2* mutant, compared to wild-type embryos. *tbg-1(RNAi)* slightly reduces EBP-2 signals at *atx-2* mutant centrosomes whereas *tbg-1(RNAi)* drastically decreases EBP-2 signals at wild-type centrosomes (n>30), illustrating the effects on MT nucleation by *tbg-1(RNAi)*. In contrast, *klp-7(RNAi)* increases centrosomal EBP-2 signals in both wild-type and mutant embryos. KLP-7 knockdown also affects cytoplasmic MT growth patterns. Magnified regions illustrate that while MTs emanate from the centrosome and reach the cortex in the wild-type embryo, EBP-2 tracks in mutant embryos appear randomly oriented. (B) Kymograph analysis in *klp-7(RNAi)* vs control treated wild-type and *atx*-2 mutants: Levels of EBP-2-GFP signals relative to control RNAi are presented for wild-type and *atx-2* mutants, respectively. While *klp-7(RNAi)* increases MT nucleation by over 2-fold in wild-type (\*\**p*=0.004), relative increase in MT nucleation by *klp-7(RNAi)* is much lower (1.2-fold) in control *atx-2* mutants (*p*=0.6). This difference in MT nucleation by *klp-7(RNAi)* suggests a limited supply of free tubulins available for MT nucleation by mutant centrosomes. The same trends are observed for the midpoint measurement. Interestingly, relative increase in the number of EBP-2 foci near the cortex is higher (*p*=0.1) in *atx-2* mutants, unlikely due to lengthened MTs growing out from centrosomes. (C) Embryos expressing GFPγ-Tubulin reveal that in *klp-7(RNAi)* in the wild-type embryo, MT nucleation is increased and MTs appear to grow longer, originating from centrosomes. However, while more MTs (based on the GFPγ-Tubulin signal) are found in the cytoplasm distant from the centrosome in *klp-7(RNAi)*; *atx-2* embryos compared to *L4440*; *atx-2* embryos, these MTs seem to be disorganized and appear disconnected from the centrosome, which might explain the increase we observe in EBP-2 signal by cortex kymograph analysis, as opposed to an increase in MT growth. These abnormal patterns of astral MTs can be reminiscent of the result previously observed by Srayko et al., (2005), in which freely moving MTs are detached from the centrosome owing to an excessive MT nucleation by KLP-7 depletion. Embryos expressing GFPγ-Tubulin reveal that *tbg-1(RNAi)* significantly decreases the number of MTs growing out from wild-type centrosomes, while *tbg-1(RNAi)*-treated mutant centrosomes still exhibit a decent level of MT nucleation although centrosomal MT signal is reduced compared to control. (A, C) Boxes are magnified 2-fold. (D) *klp-7(RNAi)* in *atx-2* mutants leads to a synergistic increase in embryonic lethality. *klp-7(RNAi); atx-2(ne4297)* animals produced ~90% of embryonic lethality compared to 30% in *atx-2(ne4297)* and only 6% in *klp-7(RNAi)* animals, suggesting that a small increase in the already aggravated MT nucleation and freely moving detached MTs by KLP-7 depletion negatively impacted embryogenesis. This increase in embryonic lethality might be due to enhanced cytokinesis failure resulting from exacerbated MT nucleation. *** *p*<0.001 (E) Knocking down both KLP-7 and ATX-2, indeed, leads to a significant increase in cytokinesis failure compared to single depletion. ** *p*<0.01 Each dot on the graph represents a centrosome. Horizontal bars indicate average values. Error bars are SD. Bar, 5 μm.

**S1 Movie. ATX-2 is required for proper cell division.** Embryos expressing GFP-γ-Tubulin, mCherry-γ-Tubulin and mCherry-H2B. The *atx-2(RNAi)* embryo contains extra DNA, presumably resulting from polar body extrusion failure during meiosis (see **S3 Movie**). Before pronuclei meet, the spindle starts to form only around the sperm pronucleus, followed by later incorporation of the maternal DNA into the spindle. The *atx-2(RNAi)* embryo displays abnormal spindle positioning, improperly positioned axis of the metaphase plate, and lagging DNA at first anaphase. The *atx-2(RNAi)* embryo duplicates centrosomes properly but fails to complete cytokinesis (see **S2 Movie**), forming a tetrapolar spindle at the second mitosis. Note that in the *atx-2(RNAi)* embryo, centrosomes labeled with mCherry-γ-Tubulin is significantly larger than in the control. Also note shorter metaphase spindle and cell cycle delay in *atx-2(RNAi)* embryo compared to the control. Each frame is equal to 1 min of elapsed time (t = 0 at metaphase I; 5 fps). Bar, 5 μm.

**S2 Movie. ATX-2 is required for normal cytokinesis.** Embryos expressing GFP-MOE marking Actin. Arrows indicate initial position of cytokinetic cleavage furrow. In the control embryo, the cleavage furrow starts to form on both dorsal and ventral (DV) sides of the embryo symmetrically, and extend toward the center of the embryo, then meet at the center to complete cytokinesis. In the *atx-2(RNAi)* embryo, the cleavage furrow only forms on one side of the embryo, and progresses to the center of the cell but is stalled at two-thirds of DV axis, resulting in cytokinesis failure. In the *atx-2(RNAi)* embryo, cortical distribution of actin appears disorganized and scattered. Note that in the *atx-2(RNAi)* embryo, position of the cleavage furrow is not sustained throughout cytokinesis. Movies are Z-projections of the middle 2-3μm of the embryo. Each time frame is equal to 30 sec of elapsed time. (t = 0 at first indication of cleavage furrow formation; 5 fps). Bar, 5 μm.

**S3 Movie. The *atx-2* mutant embryo fails to extrude polar bodies, resulting in extra DNA.** Control and *atx-2(RNAi)* embryos expressing GFP-α-Tubulin, mCherry-γ-Tubulin and mCherry-H2B. While the control embryo displays successful polar body extrusion, in the *atx-2(RNAi)* embryo the extrusion of the second polar body fails (arrow at t=12), leading to extra DNA associated with the maternal pronucleus. Note cell cycle delay and the orientation of meiotic spindle in the *atx-2(RNAi)* embryo - the meiotic spindle aligns 90°to the cell cortex throughout the meiosis II. However, in the control embryo the meiotic spindle aligns parallel to the cortex until anaphase and rotates 90°just before polar extrusion (Yang et al., 2003). Each frame is equal to 1 min of elapsed time (t = 0 at metaphase of meiosis II; 5 fps). Bar, 5 αm.

**S4 Movie. ATX-2-GFP localizes to cytoplasm and punctate cytoplasmic foci.** A transgenic strain expressing *atx-2-gfp-3x flag* (Sarov et al., 2012). ATX-2-GFP localization is consistent with endogenous ATX-2 detected by anti-ATX-2. Each frame is equal to 2 min of elapsed time; 3 fps. Bar, 5 μm.

**S5 Movie. *atx-2(RNAi)* results in increased levels of ZYG-1 at the centrosome.** Embryos expressing GFP-ZYG-1-C-terminus (Peters et al., 2010). Throughout the cell cycle from pronuclear migration to first anaphase, centriolar and PCM-like signals of GFP-ZYG-1 are much more intense in the *atx-2(RNAi)* embryo. Each frame is equal to 1 min of elapsed time (t = 0 at metaphase I; 5 fps. Bar, 5 μm.

**S6 Movie. *atx-2(ne4297)* embryos exhibit elevated levels of centrosomal ZYG-1.** Embryos expressing GFP-ZYG-1-C-terminus (Peters et al., 2010). The *atx-2* mutant embryo shows the similar pattern to *atx-2(RNAi)* embryos: *atx-2* mutant embryos display increased levels of centriolar and PCM-like GFP-ZYG-1 throughout the first cell cycle. Each frame is equal to 1 min of elapsed time (t = 0 at first metaphase; 5 fps). Bar, 5 μm.

**S7 Movie. *atx-2* mutant embryos exhibit abnormal microtubule dynamics.** Embryos expressing EBP-2-GFP that marks microtubule plus ends. More microtubules emanates from *atx-2* mutant centrosomes, compared to the wild-type. However, only few of the microtubules that grow out from the mutant centrosome continue to grow out and reach the cortex. Note that the *atx-2* embryo exhibits increased levels of centrosomal EBP-2 signal and shorter mitotic spindles, compared to the wild-type. Movies are acquired at 500 msec intervals over a 10 sec-period; 6 fps Bar, 5 μm.

**S1 Table.** List of *C. elegans s*trains in this study.

| Name | Genotype | Origin |
| --- | --- | --- |
| N2 |  | CGC |
| WM210 | *atx-2(ne4297) III* | Mello/Shirayama Lab |
| OC14 | *zyg-1(it25) II* | Kemphues et al. (1988). *Genetics* **120**: 977-986 |
| OC133 | *szy-20(bs52) II* | Kemp et al. (2007). *Genetics* **176**(1):95-113 |
| OC196 | *szy-20(tm1997) II* | Song et al., (2008) *Dev. Cell* **15**(6): 901-912 |
| OC341 | *unc-119(ed3) III; bsIs8[pMS5.1:unc-119(+)] pie-1-gfp-zyg-1C-terminus]* | Peters et al. (2010). *J. Cell Sci.* **123**(5):795-805 |
| TH66 | *unc-119(ed3) III; [pie-1::ebp-2::GFP]* | Srayko et al. (2005). *Dev. Cell* **9**(2): 223-236 |
| SA250 | *tjIs54[pie-1p::GFP::tbb-2 + pie-1p::2xmCherry::tbG-1 + unc-119(+)]. tjIs57[pie-1p::mCherry::his-48 + unc-119(+)]* | Toya et al. (2010). *Methods Cell Biol.* **97**:359-372 |
| JH2338 | *unc-119(ed3) III; axIs1489[pCG61]* | Gallo et al. (2008) *Developmental Biology* **323**(1):76-87 |
| MAS15 | *unc-119(ed3); [pie-1::unc-119(+)] + pie-1::gfp::klp-7]* | Han et al., (2015) *PLoSOne* **10**(7):e0132593 |
| JA1334 | *wels11[unc-119(+)] + TAC-1::GFP]* | Le Bot et al. (2003) *Current Biology* **13**: 1499-1505 |
| MTU 1 | *unc-119(ed3); [unc-119(+)] + szy-20 ::gfp::3xflag]* | This study; Sarov et al., (2012) *Cell* **150**(4):855-866 |
| MTU 2 | *unc-119(ed3); [unc-119(+)] + atx-2 ::gfp::3xflag]* | This study; Sarov et al., (2012) *Cell* **150**(4):855-866 |
| MTU 3 | *unc-119(ed3); [unc-119(+)] + rpn-12 ::gfp::3xflag]* | This Study; Sarov et al., (2012) *Cell* **150**(4):855-866 |
| MTU 4 | *HA::zyg-1(II)* | This Study (**S2 Table**); Dickinson et al., (2015) *Genetics* **120**: 1035-1075; Paix et al., (2015) *Genetics* **201**: 47-54. |
| AZ244 | *unc-119(ed3); [pie-1::unc-119(+)] + pie-1::gfp::tubulin]* | Praitis et al., (2001) *Genetics* **157**(3):1217-26 |
| TH27 | *unc-119(ed3); [pie-1p::gfp::tbG-1 + unc-119(+)]* | Hannak et al., (2002) *J. Cell Biol.* **157**:591-602 |

S2 Table. CRISPR/Cas9-mediated generation of transgenic strains.

S1 Table. List of *C. elegans s*trains in this study

S2 Table. CRISPR/Cas9-mediated generation of transgenic strains Using the CRISPR/Cas-9 method (Dickinson et al., 2013; Paix et al., 2015), we generated a strain expressing HA tagged ZYG-1 at the endogenous level from the N-terminus of the native genomic locus. Cas9/sgRNA plasmids were generated for both *dpy-10* and *zyg-1* using the plasmid pDD162 (Addgene) following the protocol (Dickinson et al., 2015). Single-stranded oligonucleotide (ssODN) homologous repair templates (HRTs) were designed for both *dpy-10(cn64)* (Arribere et al., 2014; Paix et al 2015) and HA::*zyg-1*(codon optimized, IDT). Injections were carried out as described in Dickinson et al., 2015 using the following injection mix (final volume = 20 μL, stock solutions in brackets):

|  |  |
| --- | --- |
| *dpy-10* HRT [500 ng/μL]: | 0.55 μL |
| *zyg-1* HRT [2 μg/μL]: | 1.1 μL |
| *dpy-10* sgRNA plasmid [400 ng/μL]: | 2.5 μL |
| *zyg-1* sgRNA plasmid [400 ng/μL]: | 2.5 μL |
| Sterile ddWater: | 13.35 μL |

Screening was carried out as described by Paix et al., 2015 - jackpot roller/dumpy broods were identified and screened for HA insertion by single-worm PCR and restriction digestion with HpyCH4IV (NEB).

sgRNA targets used in this study (PAM domain omitted):

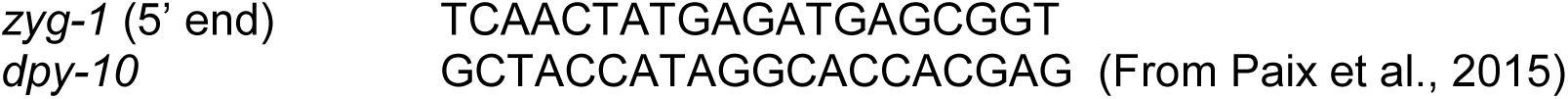

**ssODNs used in this study:**

*zyg-1* (HA sequence underlined)

tatttttagcgcaccaagtgttgatcaactatgagATGTACCCGTATGACGTGCCTGATTACGCTatgagcggtgggaagagtggttcaagattgagtgtagg

*dpy-10* (From Arribere et al., 2014) CACTTGAACTTCAATACGGCAAGATGAGAATGACTGGAAACCGTACCGCATGCGGTGCCTATGGTAGCGGAGCTTCACATGGCTTCAGACCAACAGCCTAT

**Mutagenic primers for Cas9/sgRNA plasmid generation:**

*zyg-1* sgRNA mutagenic primer:

TCAACTATGAGATGAGCGGT GTTTTAGAGCTAGAAATAGCAAGT

dpy-10 sgRNA mutagenic primer:

GCTACCATAGGCACCACGAG GTTTTAGAGCTAGAAATAGCAAGT

**HA-*zyg-1* primers for single-worm PCR:**

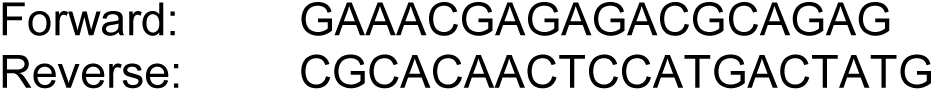

**Worms generated from this study:** MTU 4 (HA::*zyg-1*)

